# Reevaluating the Concept of Aging: Long-Term Stress Adaptation as a Key Factor in Yeast Aging

**DOI:** 10.1101/2023.11.03.565426

**Authors:** Yanzhuo Kong, Damola Adejoro, Christopher Winefield, Stephen L.W. On, Philip A. Wescombe, Arvind Subbaraj, Andrew Saunders, Venkata Chelikani

## Abstract

It has been demonstrated that short-term stress can enhance cellular responses and promote longevity, whereas long-term stress shortens lifespan. Understanding the relationship between short-term and long-term stress could offer new insights into comprehending and modulating age-related diseases. In this study, we investigate this relationship using transcriptomic and metabolomic analyses in the yeast model system (*Saccharomyces cerevisiae*).

We employed three metabolic treatments: firstly, treating yeast cells with threshold levels of benzoic acid for 24 hours (Short-term [ST] Stressed Cells); secondly, treating yeast cells with threshold levels of benzoic acid for 500 hours, with sub-culturing every 24 hours (Long-term [LT] Stressed Cells); and thirdly, allowing the long-term stressed cells to grow for 16 hours without any benzoic acid (Recovered Cells).

Here, we propose that aging is an evolutionarily conserved cellular adaptation mechanism in response to long-term stress exposure. Under short-term stressed conditions, prominent lifespan-extending metabolites such as trehalose and metabolites linked to tumor suppression in humans, such as 5’-methylthioadenosine, were overexpressed. In contrast, LT Stressed Cells activated genes such as those responsible for epigenetic regulatory enzymes that govern the aging process, and secondary stress response genes, such as heat shock proteins (HSPs) which are associated with adaptation to cell damage but also often associated with aged cells. Chronological lifespan experiments showed that LT stressed cells lived a shorter lifespan compared to ST Stressed Cells. This suggests that the markers of aging (eg. HSPs, certain epigenetic regulators) are expressed in response to long-term stress to enable cell survival but have the long-term effect of reducing lifespan. In support of this hypothesis, we also show that genes exclusively activated in ST Stressed Cells are conserved solely in eukaryotes, while those significantly expressed in LT Stressed Cells (aging related) exhibit high conservation across all domains of life, with a majority having originated from bacteria hinting at the potential evolutionary benefit of aging.

## Introduction

What constitutes the fundamental and most basic aspect of the aging process?^1^ The formerly comforting notion that aging was a natural process involving older individuals gracefully making way for the younger generation for the benefit of the species was debunked by evolutionary biologists in the latter part of the twentieth century. They demonstrated that natural selection typically prioritizes individual reproduction over the well-being of the species, rendering the idea that aging evolved to limit older individuals from reproducing for the benefit of the younger generation theoretically unsound^2,3^. Instead, it is now widely accepted that aging likely emerged later in biological history, and its precise origins and timing are central questions in evolutionary biology. Recent discoveries, such as aging in bacteria, suggest that aging predates the appearance of eukaryotic organisms, tracing its roots back to basic single-celled life forms^4,5^. No genes are known to have evolved specifically to cause damage and aging^3^. Is there another potential evolutionary benefit to aging? Although natural selection on its own may not convincingly account for the development of aging, as it often leads to the demise of most individuals, it is clear that genes play a substantial role in the processes of aging and longevity. It is probable that the genes associated with aging and longevity are part of stress response systems^6^. Nutrient and stress receptors play a role in prolonging lifespans triggered by a variety of environmental and physiological cues^7^. In-depth research has highlighted that encountering short bouts of stress can enhance the way cells respond to stress, a phenomenon referred to as “hormetic stress.” This kind of stress contributes to extended lifespans by boosting the activities of molecular chaperones and other defense mechanisms. On the contrary, enduring periods of stress can overwhelm the body’s compensatory responses, resulting in what’s known as “toxic stress,” ultimately decreasing lifespan^8,9^. According to the hyperfunction theory, as cells age, they undergo notable transformations and deficiencies at multiple levels, indicative of their adaptation to a range of stressors^10,11^. A recent research study by Poganik et al., reveals that biological aging could undergo temporary acceleration during the periods of stress. However, this acceleration is temporary, and after the stress subsides, the biological age tends to revert back to its original baseline^12^. All of this information prompts the question of whether aging might potentially function as an adaptation mechanism to prolonged stress, rather than being solely a consequence of extended stress exposure. In this context, we propose that the process of adapting to long-term stress itself constitutes aging, and aging reverses once prolonged stress is removed or in other words “aging” is in fact long-term stress adaptation, as shown in Figure 2. The figure describes the observed association between the duration of stress and either the rate of aging or the rate of recovery, with one showing a positive correlation and the other showing the opposite. If this new theory proves to be accurate, it could offer a solution to the longstanding question regarding the evolutionary purpose of aging. Gaining insights into the impacts of both short-term and long-term stress, as well as the recuperation of cells from stress, can provide valuable understanding of the aging process. Subsequently, this knowledge can pave the way for strategies aimed at addressing and mitigating the effects of aging.

The budding yeast, *Saccharomyces cerevisiae* (yeast), has served as a fundamental model organism for investigating cellular aging^13^. In particular, we believe yeast is an excellent model system to study the difference between short-term stress and long-term stress. Historically, studies focused on yeast aging were restricted to a narrow selection of genes and relied on biased research approaches. These methodologies primarily involved the evaluation of specific genes based on existing knowledge or assumptions, alongside the exploration of additional traits such as stress resistance or alterations in gene expression associated with age-related changes, which could potentially be linked to longevity^14^. While single-gene investigations in model organisms have previously provided valuable insights into the aging phenomenon^15^, we now acknowledge the significance of conducting comprehensive genome-wide inquiries into aging processes. This shift is driven by the realization that a thorough understanding of aging requires a holistic approach, as genes are intricately interconnected, and multiple homologous genes may coexist in core metabolic processes like aging. The chronological lifespan (CLS) of a yeast, is determined by assessing the duration for which cells remain viable during the late post-diauxic and stationary phases after exponential growth. Research on yeast CLS, which serves as a model for understanding post-mitotic cellular aging in more complex organisms, has been instrumental in discovering shared pathways and mechanisms that play a role in regulating lifespan^16,17^.

In this present study, we utilized benzoic acid, a widely used food preservative and antifungal agent, at sub-lethal threshold levels to establish three distinct metabolic groups. The first group involved treating yeast cells with 10mM benzoic acid for 24 hours (Short-term/ST Stressed Cells). The second group encompassed treating yeast cells with 10mM benzoic acid for 500 hours, with sub-culturing every 24 hours (Long-term/LT Stressed Cells). The third group consisted of LT Stressed Cells subsequently grown for 16 hours without any benzoic acid, allowing for recovery (Recovered Cells). In this study, we regarded a shorter lifespan and fewer cells as indicative of aging.

We performed transcriptomic and metabolomic investigations, specifically emphasizing genes and metabolites that exhibited significant expression changes in cells subjected to long-term stress (LT stressed cells). These included epigenetic regulators and secondary stress response genes, which are also recognized as key markers of the aging process^18–20^. Additionally, we conducted a phylogenetic analysis of these genes to trace their evolutionary origins.

Our findings reveal notable differences in the transcriptomic and metabolomic profiles between LT Stressed Cells and ST Stressed Cells. The chronological lifespan of LT Stressed Cells proved to be shorter than that of ST Stressed Cells, with several age-related biomarkers being prominently expressed in the former. Several genes were specifically expressed in LT Stressed Cells, suggesting the existence of a potentially distinct mechanism, separate from the primary stress response, contributing to aging in these cells. Furthermore, our phylogenetic analysis of genes specifically expressed in LT Stressed Cells indicates high conservation, primarily from bacterial origins. This aging process observed on LT Stressed Cells seems to operate independently yet in harmony with stress response pathways, potentially offering valuable support for cell survival. Recovered Cells exhibited similarities to Control Cells, with several lifespan-extension genes and metabolites activated in line with existing literature. Additionally, both Stressed and Recovered Cells produced several beneficial metabolites, hinting at potential biotechnological applications stemming from these cell states.

## Results and Discussion

### Stress Responses

#### Metabolic profiles

The body’s longevity response to dietary restrictions and stress has been shown to be actively governed by pathways that sense stress and nutrients^7,21,22^. In model organisms, restricting methionine has been shown to extend lifespan and delay the onset of age-related diseases. Previously, Ogawa et al. (2016 & 2022) observed that the metabolite S-adenosyl-L-homocysteine (SAH) activates adenosine monophosphate (AMP)-activated protein kinase (AMPK) in yeast, leading to extended lifespan. Their recent research demonstrates that SAH supplementation can decrease methionine levels and reproduce many of the physiological and molecular effects associated with methionine restriction in *Caenorhabditis elegans*. Treating with SAH, prolongs lifespan by activating AMPK and providing similar benefits to methionine restriction^23,24^.

To further explore the impact of short term and long term stress on yeast, we utilized benzoic acid, a widely used food preservative and antifungal agent, at sub-lethal threshold levels to establish three distinct metabolic groups. The first group involved treating yeast cells with 10mM benzoic acid for 24 hours (Short-term/ST Stressed Cells). The second group encompassed treating yeast cells with 10mM benzoic acid for 500 hours, with sub-culturing every 24 hours (Long-term/LT Stressed Cells). The third group consisted of LT Stressed Cells subsequently grown for 16 hours without any benzoic acid, allowing for recovery (Recovered Cells). RNA transcriptomics and metabolomics were then used to identify changes between the three groups in gene expression and metabolite production respectively.

The major findings of this study are presented in Figure 1, which summarises both transcriptomic and metabolomic results in terms of the up- and down-regulation of key genes and metabolites in all treated yeast cells compared to the Control Cells. One clear finding from the metabolomic analysis was increased production of SAH, which is shown in both Figure 4 (metabolite 12 [negative charge mode], and metabolites 40 & 41 [positive charge mode]) and Figure 6, in Recovered Cells compared to Control Cells. Both figures depict the metabolic profiles in all types of yeast cells, as detected by LC-MS/MS analysis, with Figure 4 presented as a circular heatmap and Figure 6 emphasising four key metabolites. Interestingly, the SAH production was significantly down-regulated in LT Stressed Cells (opposite of Recovered Cells). It is highly likely that activation of SAH production is necessary for cell survival after long-term stress exposure. Our findings align with the recent study by Poganik et al. (2023) which proposes that aging is a temporary process heightened by stress and subsequently restored during recovery. The up-regulation of SAH production play a pivotal role in aiding cell recovery following exposure to stress^12^. Therefore, our finding indicate that SAH levels in a cell could potentially serve as a biomarker for recovery (Figure 2).

**Figure 1.**
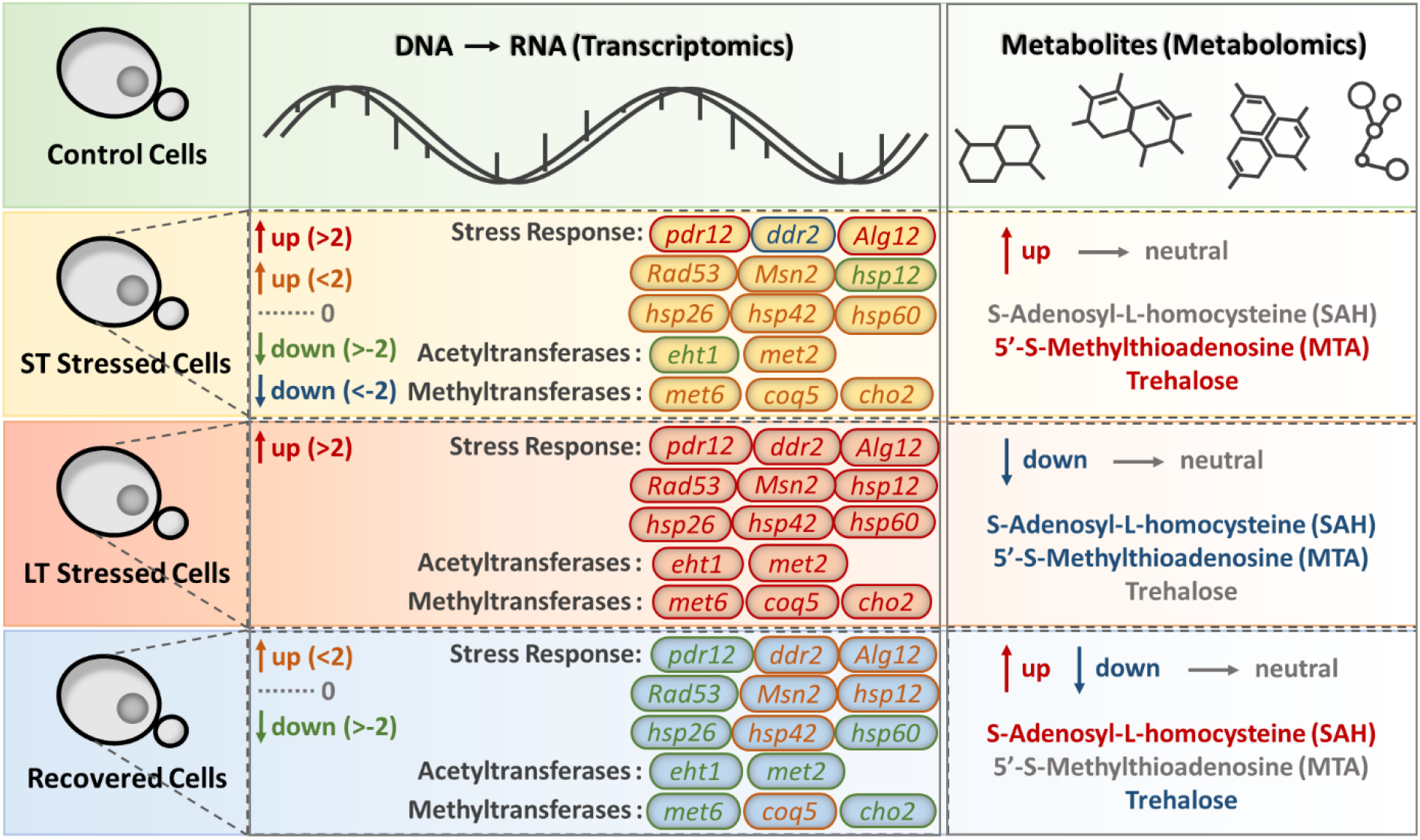
Transcriptomic and metabolomic changes in *Saccharomyces cerevisiae* key genes and metabolites in response to epigenetic modification. The labels for up- (red and orange) and down-regulation (green and blue) in the transcriptomic section are determined using log2 fold change values obtained from the results of differential gene expression analysis. **Control Cells:** wild type *S. cerevisiae*; **ST Stressed Cells:** short-term stressed cells, *S. cerevisiae* firstly exposed to 10 mM benzoic acid for 16 hrs; **LT Stressed Cells:** long-term stressed cells, *S. cerevisiae* exposed to 10 mM benzoic acid for 500 hrs (24 hrs/sub-culture); **Recovered Cells:** *S. cerevisiae* exposed to 10 mM benzoic acid for 500 hrs (24 hrs/sub-culture), followed by growing in regular YPD broth for 16 hrs.

**Figure 2.**
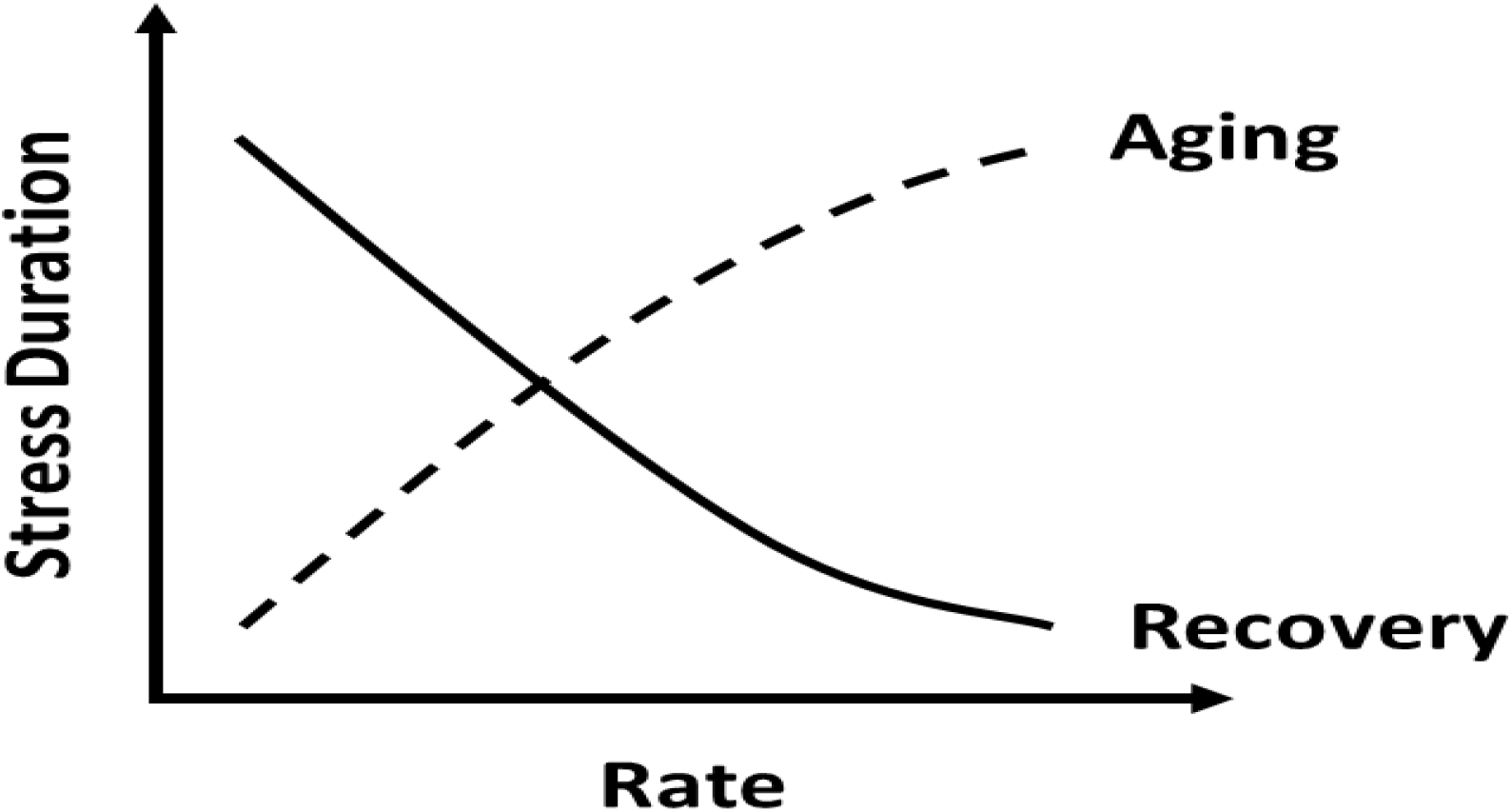
The proposed relationships between stress response and aging/recovery highlighting the possibility that aging is actually the result of a cells survival mechanism in response to stress.

Yeast cell viability and morphology were evaluated using the chronological lifespan experiment (figure 3). This showed that LT Stressed Cells have a relatively shorter lifespan than ST Stressed Cells, but that they recovered immediately once the stress was removed from the environment. The morphology of ST and LT Stressed Cells was found to be similar, however, the Recovered Cells exhibited an improved morphological appearance compared to the Control Cells (Figure 3c). The observation that the overproduction of SAH did not significantly affect lifespan, contradicts the existing evidence that SAH supplementation/overproduction alone can extend lifespan^24^. Nonetheless, in our work, SAH does appear to play a role in facilitating recovery from long-term stress exposure and may contribute to slight improvements in cell morphology and a modest increase in chronological lifespan among recovered cells following the removal of stress.

**Figure 3.**
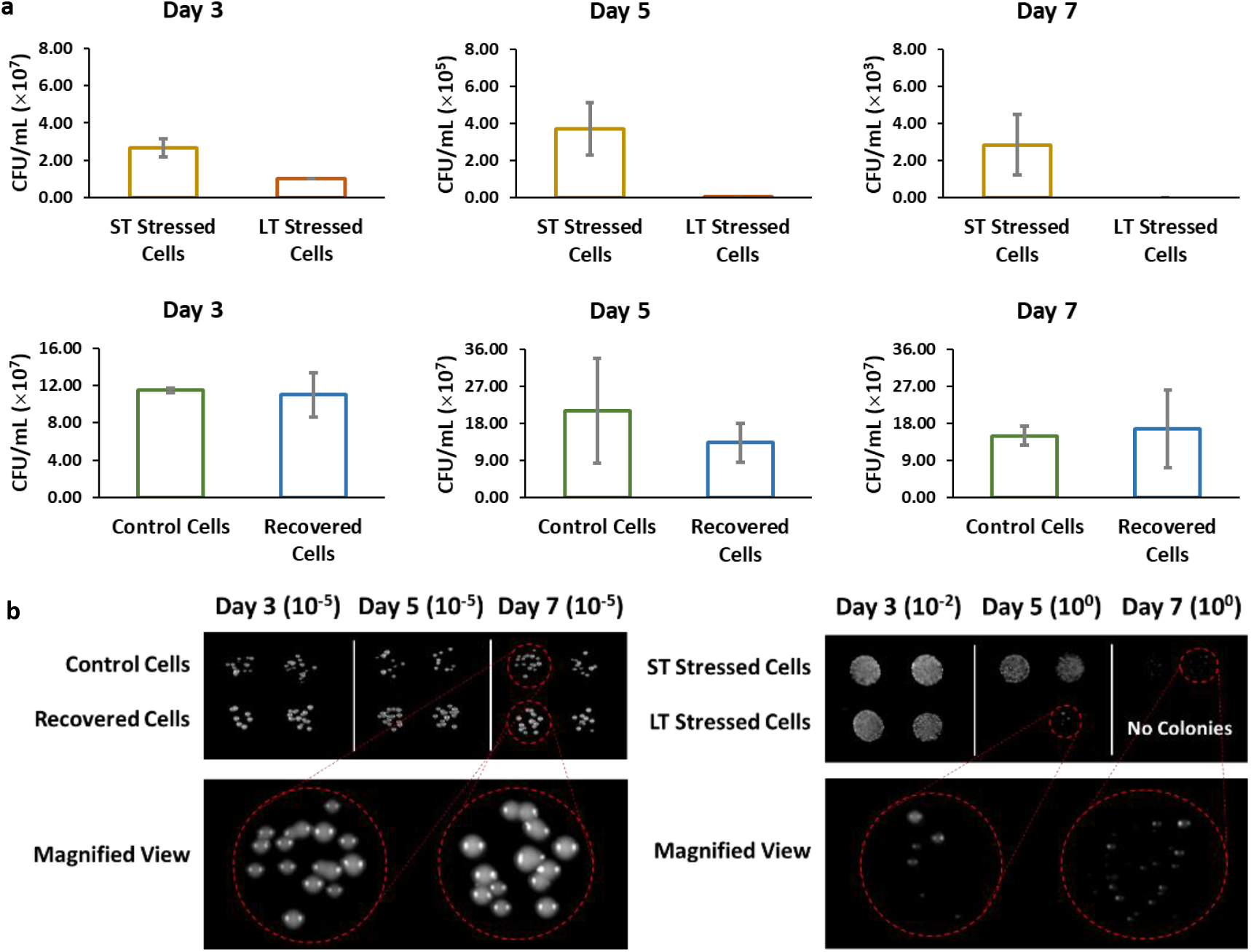
Chronological aging experiment of *S. cerevisiae* exposed to different stress durations. (**a**) cell viability (*p* < 0.05 between ST Stressed Cells and LT Stressed Cells, *p* > 0.05 between Control Cells and Recovered Cells) and (**b**) morphology on day 3, 5 and 7. The initial colony forming units (CFU/mL) of each sample were uniformed on day 0, at 10^6^ CFU/mL. For morphology observation, 10 µL of serially diluted *S. cerevisiae* cells were plated on YPD agar plates which were then incubated at 37 ℃ for 24 hrs.

Interestingly short-term stress did not exert a significant impact on SAH production despite increasing the prevalence of most other metabolites measured (Figure 4,6). Included in the metabolites which were over produced in the presence for short term stress were trehalose and 5’-S-methylthioadenosine (MTA). Trehalose, a well-established metabolite, is recognized for its ability to enhance lifespan^25–27^. MTA is a nucleoside derived from S-adenosyl methionine (SAM)^28,29^, known for its beneficial properties, including the suppression of tumours^30^. It has been previously demonstrated that enhancing SAM synthesis can lead to an extended lifespan. SAM is generated from methionine through the action of methionine-adenosyltransferases (MATs) and functions as a methyl group donor for various biological processes. In the course of this metabolic pathway, SAH is formed in a reversible reaction. It is worth noting that SAH has been shown to act as an inhibitor of methyltransferases^24,31,32^. Significantly, both SAH and MTA were downregulated in LT Stressed Cells. These findings lend support to our hypothesis that short-term stress has the potential to activate mechanisms associated with extending lifespan, aligning with existing literature ^8,33^. Conversely, long-term stress appears to initiate processes linked to aging, as evidenced by the downregulation of SAH production in our study.

#### Transcriptional profiles

The gene expression profiles of treated yeast cells, compared to control cells, are presented in Figure 5. This figure includes four categories of genes based on their roles and functions, which are stress response, autophagy process, methyltransferases, and acetyltransferases. The cellular defence mechanisms that guard against various threats, such as high temperatures, oxidative stress, or osmotic stress, comprise multiple layers of protection. The initial line of defence comprises small molecules with low molecular weight, such as trehalose, activation of essential protein repair systems and chaperones like *pdr12*^34^ and *pdr5*^35^ to survive chemical stress, and upregulation of genes like *tos*4^36^ to promote gene expression homeostasis. These components are vital for ensuring immediate survival in adverse conditions. In this study, trehalose which is a widely known lifespan extension compound was significantly overproduced in ST Stressed Cells. This observation supports their dual role in both the first line of defence and also in lifespan extension in the absence of external stresses. The continuation of stress exposure then activates secondary processes such as upregulating the genes responsible for producing protective factor, this is the point where we believe cells start the process of aging and activate secondary stress response mechanisms for cell survival such as heat-shock proteins. Heat shock proteins (HSPs) are highly prevalent and evolutionarily preserved protein groups found in both prokaryotic and eukaryotic life forms. They play a crucial role in preserving the stability of cellular proteins and shielding cells from various stressors. The classification of HSP protein families is primarily determined by their molecular sizes, with notable categories encompassing large HSPs, Hsp90, Hsp70, Hsp60, Hsp40, and small HSPs, including Hsp40, Hsp60, Hsp70, Hsp90, and Hsp110^37,38^ There is compelling evidence demonstrating age-related increases in the expression of HSPs. For instance, in the nematode *Caenorhabditis elegans*, several human HSP homologs (e.g., Hsp-16.48, Hsp-43, Hsp-17, and Sip-1) were found to increase in abundance as the nematodes aged^39^. Additionally, elevated transcript levels of HSPs were associated with reduced heat shock resistance, disrupted proteostasis, and shorter lifespan in these nematodes^39^. In Drosophila, the aging process was observed to lead to an upregulation of Hspb8 and HspA^40,41^. The expression levels of these HSPs were predictive of the lifespan of adult flies under normal aging conditions and when subjected to heat or oxidative stress. The alteration of HSP levels in relation to advanced age remains a subject of debate^42^.

**Figure 4.**
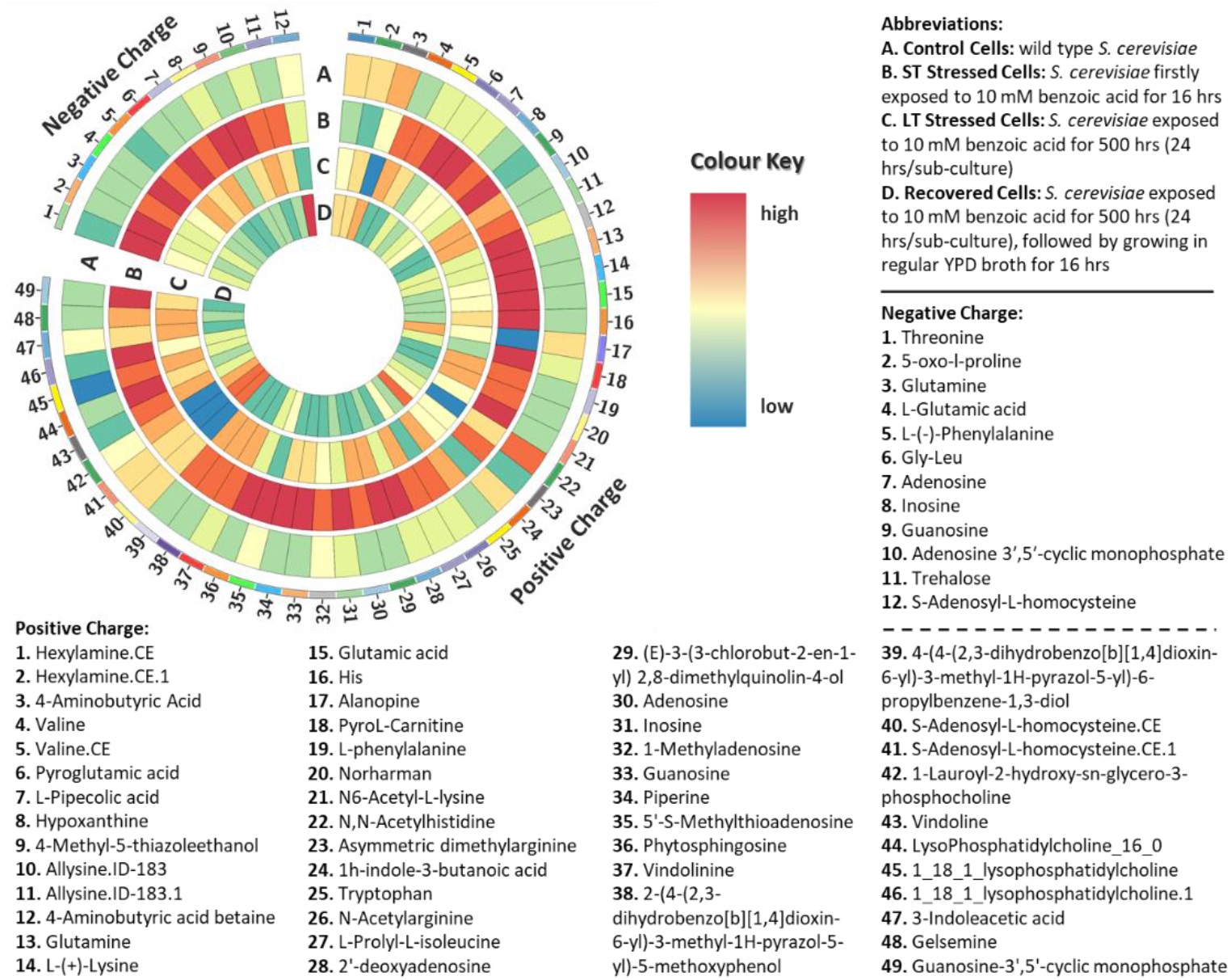
Circos plot of known metabolite concentrations detected by LC-MS/MS metabolomic analysis in control and treated *S cerevisiae* cells, including both positive and negative charge modes. Colour key indicates concentration of metabolite detected high (red), low (blue) of each metabolite produced by different *S. cerevisiae* cell types.

Hence, there must be an overarching mechanism capable of detecting and reacting to various stress forms. Initially, a regulatory element responsive to multiple stress conditions was identified as an Heat shock factor-independent heat shock element in the promoters of *ctt1*^43^ and *ddr2*^44^. This element, was termed the stress response element (STRE) and has demonstrated its ability to mediate transcription induced by diverse types of stress^44–48^. In line with its role in promoting general stress resistance, STRE has been identified as the controller of stress-inducible transcription in genes responsible for protective functions. The roster of genes under the influence of STRE includes *ctt1*, *ddr2*, *hsp12*, *tps2*, *gsy2*, and *gph1*^47,48^. The mechanisms responsible for detecting and transducing various stress signals via STREs remain largely undisclosed.

We show that long-term stress activates STRE and activates all the genes (*ctt1*, *ddr2*, *hsp12*, *tps2*, *gsy2*, and *gph1*) under the influence of STRE (Figure 5b). Previous research has shown that *msn2*, *msn4* and *gis1*^48–51^ are essential for the activation of numerous yeast genes like *ctt1*, *ddr2*, and *hsp12*, whose upregulation occurs via STREs and mutants deficient in *msn2* and *msn4* are known to be hypersensitive to severe stress conditions^52^. These genes encode a DNA-binding component of the stress-responsive system, and it’s probable that they function as positive transcription factors. All three *msn2*, *msn4* and *gis1* are significantly upregulated in LT Stressed Cells. We also noted a substantial rise in the levels of another HSP gene, *hsp104*. Research has indicated that *hsp104* consistently elevates its concentration as cells age. The concentration of *hsp104* followed a steady increase throughout the aging process, reaching a two-fold increase compared to young cells at the time of the final budding event^53^ . Notably, this increase continued even beyond this point, with *hsp104* concentrations rising further during the subsequent posterior G1 arrest phase at an even higher average rate^53^. Several other HSP genes, such as *hsp30*, *hsp26*, *hsp78*, *hsp42*, *hsp12* and *hsp82* are all over expressed in LT Stressed Cells (Figure 5b). No significant over expression of HSP genes were observed in ST Stressed Cells and down regulation of these HSP genes were observed in Recovered Cells. HSPs have been linked to both cancer and neurodegenerative diseases, and it is well-established that inhibitors targeting these proteins have demonstrated efficacy against these conditions^54–56^. Exploring these HSP inhibitors may hold promise as a potential approach for intervening in the aging process.

**Figure 5.**
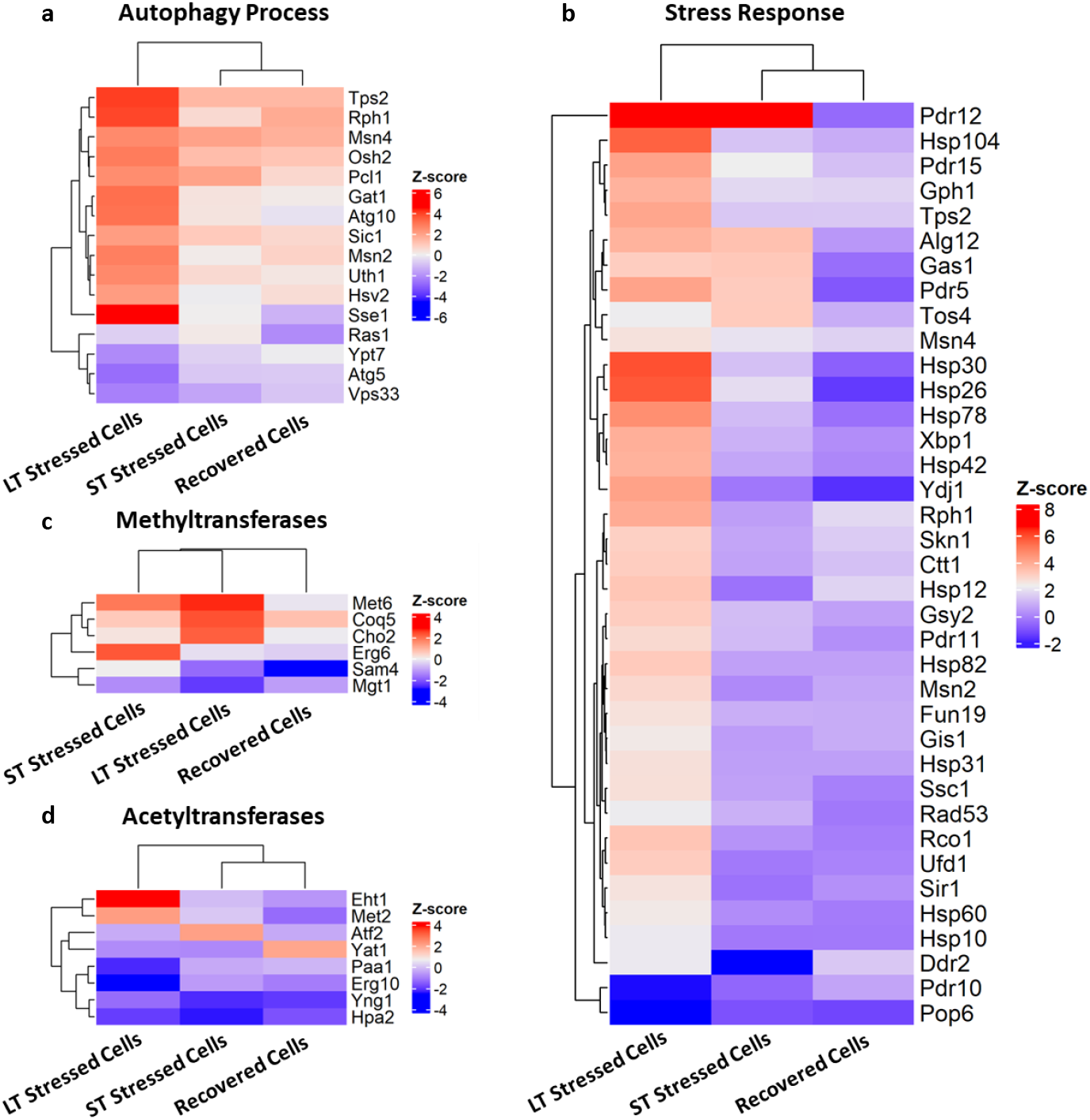
Differentially expressed genes in treated *S. cerevisiae* cells in comparison with control cells, which play an important role in: **(a)** autophagy process; **(b)** stress response; or in encoding transferases crucial for **(c)** methylation; and **(d)** acetylation. Z-score normalisation was applied during data visualization, where red and blue colour were used to represent up- and down-regulation of relevant genes.

**Figure 6.**
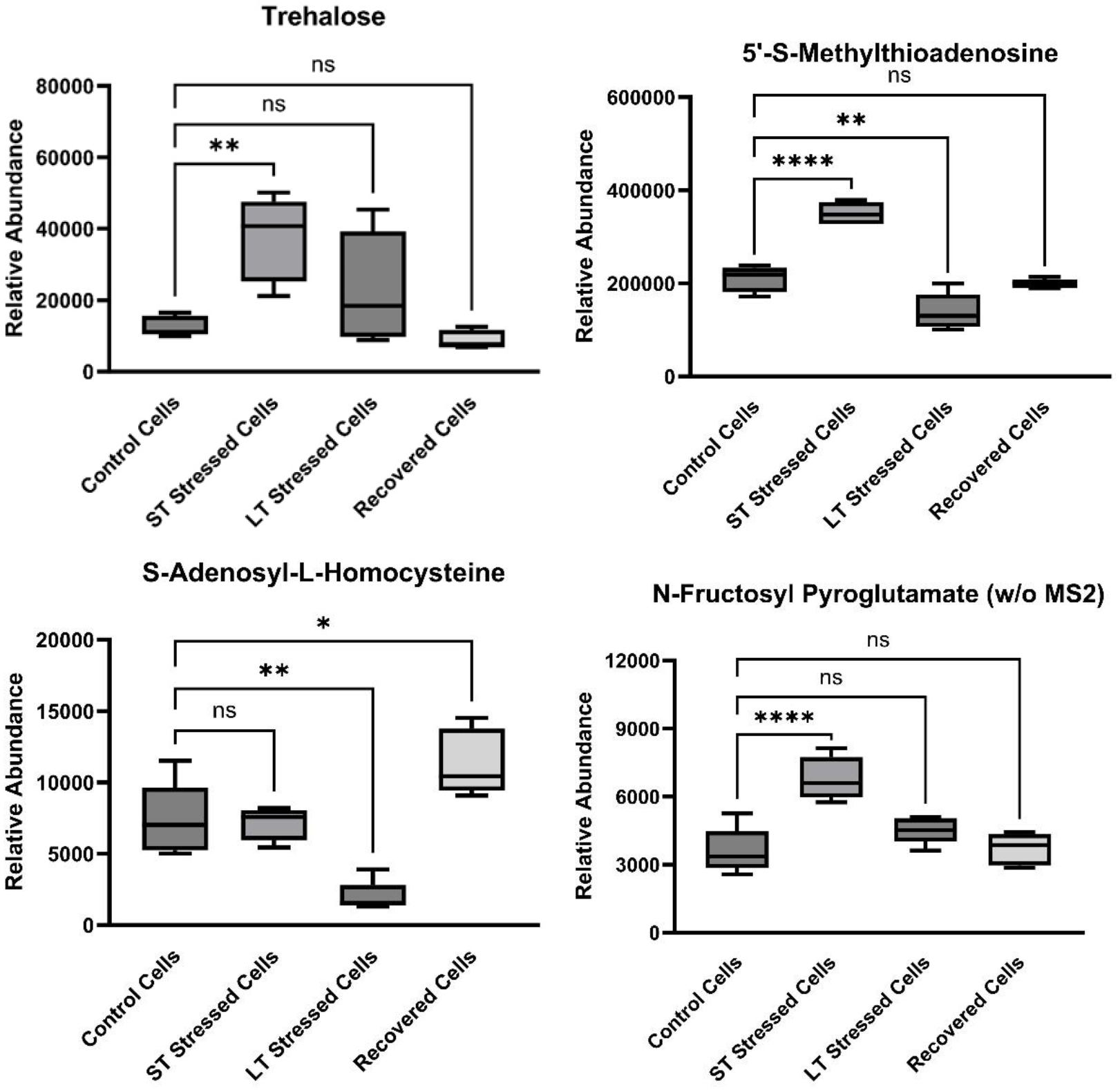
The relative abundance of representative metabolites detected by LC-MS/MS, in *S. cerevisiae* cells differentially exposed to benzoic acid. The annotation ns, indicates the relevant samples were not significantly different. Asterisks indicate the relevant samples were significantly different (* *p* < 0.05, ** *p* < 0.01, **** *p* < 0.0001).

Furthermore, Labunskyy et al. made an intriguing observation of unexpected lifespan extension in unfolded protein response (UPR) target gene deletions, such as *alg12* and *blt1*. This discovery contradicted initial expectations, as these genes are typically deemed crucial for restoring ER homeostasis and bolstering cellular protection. The authors of this study proposed that this surprising outcome could be linked to hormesis, where mild stress triggers protective mechanisms against more severe age-related stressors^57^. Interestingly, this aligns with our hypothesis that aging represents a long-term stress response and adaptation, with the deletion of these genes contributing to lifespan extension. In our investigation, we found a significant upregulation of *alg12* in both ST and LT Stressed Cells, with fold increases of 3.4 and 3.8, respectively (see Figure 5b) Indicating that increased upregulation can result in a reduced lifespan, while its deletion leads to an extension of lifespan^57^. Additionally, it’s worth noting that the mechanisms underlying lifespan extension due to UPR target gene deletion appear distinct from those associated with increased *sir2* activity or reduced mTOR signaling, which mimic the effects of dietary restriction^57^.

Our findings also revealed no significant increase in *sir2* activity or *sch9* and only a slight reduction in mTOR activity, indicating the potential existence of separate pathways for nutrient and stress sensing in the context of aging and lifespan extension. In addition, *rad53* has previously been associated with aging but had not been connected to the response to dietary restriction. It is established that *rad53* induces cell cycle arrest in response to DNA damage caused by MMS^58–60^. Importantly, in our research, we observed a significant upregulation of *rad53* expression in LT Stressed Cells (Figure 5b).

### Epigenetic Responses

Epigenetic responses form an essential component of the stress response^61^. There is substantial evidence endorsing the idea that epigenetic influences play a pivotal role in controlling the aging process. For instance, research has identified the occurrence of epigenetic changes as individuals age, known as “epigenetic drift.” Importantly, the aging rate can be directly influenced by a combination of environmental and epigenetic factors. Continuous exposure to environmental stressors throughout one’s life may lead to epigenetic alterations both at the cellular and organismal levels^20^. Chromatin-modifying enzymes employ cellular metabolites as their substrates, thus establishing a connection between metabolic pathways and the processes of epigenetic modification and gene regulation. The one-carbon cycle, mediated by enzymes like MATs, regulates gene expression through epigenetic mechanisms, with SAM and SAH acting as activators and inhibitors, respectively, of DNA and histone methylation. This metabolic control of chromatin dynamics plays a pivotal role in various biological processes^62,63^, including aging.

Research has demonstrated that inducing temporary hormetic mitochondrial stress, which includes an elevation in mitochondrial reactive oxygen species (mtROS), triggers advantageous reactions that prolong lifespan. This pathway detects mtROS differently from the response to nuclear DNA damage and, in the end, enhances longevity by deactivating the histone demethylase Rph1, particularly within subtelomeric heterochromatin region demonstrating communication pathway linking mitochondria and telomeres, which play a role in governing the aging process and lifespan^17^. In our study, we observed the activation of *rph1* in cells exposed to prolonged stress (toxic stress) over an extended period (as depicted in figure 5a), indicating it may play a role in extending lifespan during short-term stress and contributing to aging during long-term stress exposure.

Research has established that the HDA complex, categorized as a class-II histone deacetylase (HDAC), governs the aging process by influencing stress response pathways, especially those involved in DNA damage and osmotic stress response. Inhibiting HDA can lead to increased lifespan by inducing the activation of the trehalose metabolic pathway. This discovery presents intriguing possibilities for addressing age-related diseases^64^, particularly in light of the existing use of multiple HDAC inhibitors for cancer treatment^64,65^. This present study observed the activation of the components of the HDA complex in LT Stressed Cells, and this activation is likely to have had an adverse effect on longevity, potentially resulting in accelerated aging in these LT Stressed Cells (Figure 5). One example component of the HDA complex that was activated is Xbp-1. Xbp1 functions as a transcriptional repressor that remains dormant during the logarithmic growth phase, yet it becomes activated in response to a wide range of stress stimuli and plays a role in the repair of DNA double-strand breaks in yeast through Xbp1-dependent histone H4 deacetylation^66^. In quiescent cells, *xbp1* transcripts are the most abundant and have the capability to repress around 15% of all yeast genes as the cells transition into the quiescent state. It is also important Numerous studies in higher eukaryotes have revealed the induction or detection of *xbp1* expression in various types of cancer cells^67^, as well as in conditions such as stroke^68^ and neurodegenerative diseases, including Alzheimer’s disease^69^. The overexpression of *xbp1* has demonstrated certain advantageous effects in the aging of the mammalian brain^69,70^. In our research, we observed a significant increase in the expression of *xbp1* in LT Stressed Cells. This suggests that *xbp1* could serve as a potential biomarker for aging, with its expression potentially playing a role in cell survival and adaptation mechanisms during the aging process or as an integral component of the aging process itself.

The gene responsible for Rco1, which forms a homodimer in the Rpd3s histone deacetylate complex^71^, was also expressed significantly higher in LT Stressed Cells compared to ST Stressed Cells and Recovered Cells (Figure 5b). This indicates that the inhibition of HDAC leads to lifespan extension, and conversely, that activation of HDAC leads to aging in the yeast cells.

Furthermore, numerous chromatin-modifying enzymes, including methyltransferase genes like *met6*, *coq5*, *cho2*, and acetyltransferase genes such as *eht1* and *met2*, exhibited significantly higher expression levels in LT Stressed Cells as compared to both ST Stressed Cells and Recovered Cells (Figure 5). Coq5 serves as the catalyst for the sole C-methylation process involved in the formation of coenzyme Q (Q or ubiquinone) in both humans and the yeast Saccharomyces cerevisiae. Within the yeast’s Q production pathway, Coq5 is one of the eleven essential polypeptides and is known to participate in the assembly of a multi-subunit complex known as the CoQ-synthome. In the case of humans, mutations occurring in various COQ genes result in primary Q deficiency, and a reduction in the biosynthesis of coenzyme Q is linked to the development of mitochondrial, cardiovascular, kidney, and neurodegenerative disorders^72^. Previous investigations in yeast have revealed that specific point mutations located within or in close proximity to the conserved methyltransferase motifs of COQ5 result in the impairment of Coq5’s methyltransferase function. Notably, mutants carrying these specific alleles (coq5-2, coq5-5) still maintain consistent levels of Coq5 protein. When these yeast mutants with the mentioned point mutations were supplemented with ubiE, an analogue of E. coli’s COQ5, the restoration of respiration and C-methyltransferase activity was observed^73,74^.

Moreover, the introduction of human COQ5 into yeast mutants by Nguyen et al., underscores the functional resemblance between the yeast and human pathways involved in the biosynthesis of Q. This discovery holds significant implications for the diagnosis and treatment of Q10 deficiencies in patients, highlighting the potential for gaining insights across species to address this concern^72^.

Methylation-driven gene silencing is crucial in colorectal and pancreatic cancer progression. hsp90 regulates DNA methyltransferase (DNMT) enzymes. Inhibiting hsp90 with ganetespib (known inhibitor of *hsp*90) reduced DNMT1, DNMT3A, and DNMT3B levels in HT-29 and MIA PaCa-2 cells, leading to reduced DNA methylation and gene reactivation^75^. Among other instances, *hsp60* is noteworthy. Expression levels of *hsp60* are heightened in various types of cancer^54^ . Furthermore, translational modifications of *hsp60*, such as acetylation, have been observed in several cancer types^76^ . Additionally, *hsp70* chaperones are implicated in numerous diseases, including cancer, inflammation, and Alzheimer’s disease^77^. Epigenetic modifications are known to regulate the *hsp70* pathway^37^.

In this present study, both *hsp60* and *ydj1*, which play a vital role in distinguishing among *hsp70* isoforms and facilitating substrate transfer to *hsp70*, were found to be significantly expressed in cells subjected to prolonged stress. This indicates that further understanding and identifying methods for targeting HSP, could not only create opportunities for treating age-related diseases, but also potentially intervene in the aging process.

### Autophagy Response

Autophagy, a cellular process for recycling, is triggered by stressors like nutrient scarcity, viral infection, and genotoxic stress. Recent evidence points to oxidative stress, mediated by reactive oxygen species (ROS) and reactive nitrogen species (RNS), as the common signal driving autophagy. While several potential mechanisms explaining the intricate connection between oxidative stress and autophagy have been proposed, only a handful of them have been confirmed to influence autophagy. Therefore, gaining insight into the precise molecular control of autophagy by ROS and RNS, as well as the close association between metabolism and the redox state, holds significant promise for advancing anticancer treatments and the development of targeted therapies for in the future^78^. Research has demonstrated that the overexpression of *atg5* activates autophagy and prolongs the lifespan of mice while enhancing their resistance to oxidative stress. Notably, this increased tolerance to oxidative stress can be reversed by the use of any autophagy inhibitor. The initial genetic link between autophagy and aging was established in *Caenorhabditis elegans*, where the reduction of *beclin1* in *daf-2* mutants dramatically shortens their lifespan^79^. Intriguingly, in this present study, it has been observed that *atg5* is significantly downregulated in LT Stressed Cells (Figure 5a). This finding suggests that *atg5* is not only implicated in extending lifespan but also in the aging process itself, as its downregulation appears to be associated with aging. However, this observation does not align with our hypothesis, necessitating further investigation. Our proposed hypothesis suggests that aging serves as a highly conserved secondary stress response mechanism across all domains of life. It is important to note that autophagy is not conserved in prokaryotes. Additionally, another aspect requiring in-depth examination is telomere attrition, as telomerases play a pivotal role in the aging process. Yet, it’s noteworthy that these genes are not highly conserved in prokaryotes. One plausible explanation is that these characteristics serve as primary stress response mechanisms, predominantly originating from eukaryotic ancestry. Alternatively, these traits could be secondary stress responses that have evolved in eukaryotes to enhance complex cell survival.

### Phylogenetic Analysis

The research findings are intriguing, highlighting distinct patterns in gene activation in response to different stress durations. Genes exclusively activated in ST Stressed Cells are conserved solely in eukaryotes, while aging-related genes significantly expressed in LT Stressed Cells exhibit high conservation across all domains of life, with a majority having originated from bacteria. Figure 7 shows the gene cooccurrence of all regulated genes across all domains of life, and supplementary Figures 1 to 5 illustrate the maximum likelihood phylogenetic trees of five highlighted yeast genes: *coq5*, *eht1*, *sse1*, *tor1*, and *hsp60*, along with their homologs across different domains of life. Among these genes, notable candidates include secondary stress response genes such as *hsp* and the epigenetic modifying enzyme *coq5*. The presence of *hsp60* homologs in various life domains and phylogenetically distant organisms raises questions about their origin, suggesting either a common ancestral source or horizontal gene transfer between different lineages. Additionally, well-supported clades of Archaea homologs of *hsp60* further emphasize the widespread distribution of this gene. Similarly, the closest human homolog of *hsp60*, the *hspd1* gene, forms a well-supported monophyletic clade with the yeast *hsp60* gene (Supplementary Figure 5). Likewise, *coq5* homologs display a wide distribution across life domains, posing the same questions about their origin and suggesting a potential horizontal gene transfer. Notably, homologs of the yeast *coq5* gene are most commonly found in the bacterial phylum Proteobacteria. Moreover, the closest human homolog of yeast *coq5*, the *tnt1A* gene, forms a well-supported monophyletic clade with homologs from the bacterial phyla Proteobacteria and Firmicutes, hinting at a conserved genetic relationship between the yeast *coq5*, the human *tnt1A*, and analogous genes present in bacteria (Supplementary Figure 1). This conservation of stress response genes across different organisms and domains of life shed light on the evolutionary history and possible benefit of the aging mechanism.

**Figure 7.**
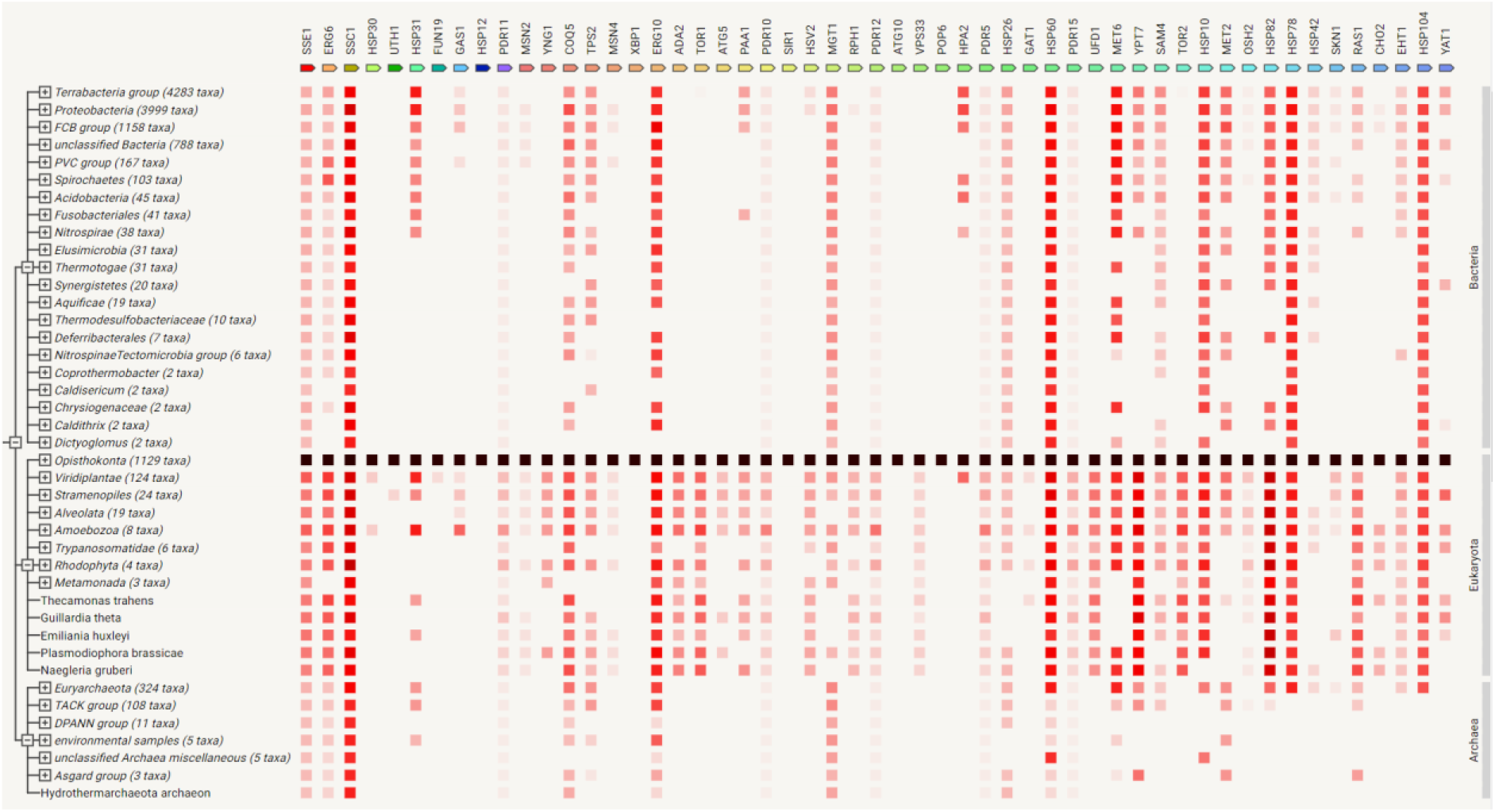
Gene cooccurrence of all regulated genes across all domains of life. The coloured square denotes, for each gene, the similarity of its best hit in a given STRING genome. The intensity of the squares denotes the degree of similarity: the more intense the square is, the more similar the gene is to its best hit. The highest intensity means a 100% sequence conservation. Black indicates the gene used for comparison.

### Biotechnology Application

Recent findings have shown that the fission yeast *Schizosaccharomyces pombe*, when cultivated at threshold levels of caffeine, developed epimutations (phenotypic changes mediated by unstable silencing rather than DNA alterations). These epimutants displayed cross-resistance to antifungal agents, suggesting that epigenetic processes can promote phenotype plasticity without modifying the genotype. In our previous research, we have shown that we can modify the yeast alcoholic fermentation process by altering cellular epigenetics through the use of epigenetic modifiers^80^. In this present study, we have demonstrated that we can alter yeast metabolism using the epigenetic modifier, benzoic acid, resulting in the differential production of 61 metabolites detected as shown in Figure 4. Among these metabolites, several have been revealed to activate the immune response, including nucleosides such as guanosine-3’,5’-cyclic monophosphate (metabolite 49, Figure 4)^81,82^ and their analogues^83,84^ which are known to act as antiviral molecules against several human viruses by acting as a chain terminator for the RNA polymerase^83^. In addition, pyroglutamic acid^85^, piperine and trehalose^86–88^ are known for their antiviral activity. SAH is also a well-known antiviral compound which was downregulated in adapted cells, it is commonly inhibited as a response to viral infection ^89^. There is the potential then, to use this approach to increase production of metabolites with biotechnology applications such as the essential amino acid L-phenylalanine which was shown to be produced in higher quantities in both ST and LT Stressed Cells^90^ (Figure 4).

## Conclusion

ST Stressed Cells activate genes such as *pdr12* and *alg12*, and metabolites such as SAM and trehalose, which provide immediate protection from stress and potential lifespan extension. In contrast, prolonged stress activates aging, which includes secondary stress response genes such as heat shock proteins, potentially causing aging as a cell stress adaptation and survival mechanism. Primary stress response genes like *pdr12* are also expressed in LT Stressed Cells which potentially help in the recovery of cells when stress is removed.

In conclusion, this study highlights the differences in short-term stress and long-term stress with respect to aging. This study provides evidence that long-term stress causes aging in *S. cerevisiae.* In addition, we hypothesize that aging could be the result of the development of a mechanism for cell survival during long-term stress episodes which provides short term benefits for cell survival when undergoing chronic long-term stress, but which leads to premature aging when the response goes on too long and subsequently shortens lifespan. The evidence presented in this present study also supports the contention that aging is reversible^12^, as demonstrated by the improved lifespan of the Recovered Cells. Furthermore, we provide evidence that primary stress response genes such as *pdr12* and *alg12*, and metabolites such as SAH and trehalose, are activated during short term stress, while secondary response genes such as *hsp12* and *hsp60* are activated during long term stress. Stress response and longevity-related genes like tor1 are remarkably preserved across eukaryotes, but not so much in prokaryotes. On the other hand, genes that are active in cells enduring prolonged stress, such as epigenetic enzymes like methyl transferase coq5, and the secondary stress response genes such hsp60, exhibit conservation across all life domains, even including prokaryotes (Figure 7). One of the most well-established concepts in biology is that all life undergoes change through natural selection. We provide evidence that indicates that the aging process can confer an evolutionary advantage during times of stress, through enhancing short-term survival advantages for a cell while having the secondary effect of reducing lifespan. Finally, the use of benzoic acid which is a known epigenetic modifier to induce stress conditions, resulted in the increased production of several beneficial metabolites. These included antiviral compounds which could have future clinical potential, and thus illustrates a possible application for using epigenetic modification as a non-GMO approach to manipulating the traits and behaviors of organisms.

Whether aging leads to adaptation to long-term stress exposure, or if aging itself is a mechanism contributing to survival in the face of prolonged stress, requires additional investigation. While it is well-known that nutrient sensing pathways and stress sensing pathways are interconnected, this study did not establish a clear connection. For instance, *tor1* was downregulated in recovered cells as anticipated, but not to a statistically significant extent, suggesting that their modes of action may be independent of each other.

## Materials and Methods

### Stress Conditions on Model Strain

A commercial yeast strain, *S. cerevisiae* EC-1118 (Lalvin EC1118™), was used as the model strain in this study. Untreated wild-type *S. cerevisiae* cells were used as Control Cells. Short-term (ST) Stressed Cells were obtained by exposing *S. cerevisiae* to 10 mM benzoic acid for 24 hrs, in comparison with Long-term (LT) Stressed Cells, which were continuously exposed to 10 mM benzoic acid for 500 hrs (24 hrs/sub-culture). Recovered Cells were LT Stressed Cells that were subsequently grown in regular Yeast Peptone Dextrose (YPD) broth for 16 hrs.

### RNA Preparation for Transcriptome Sequencing

*S. cerevisiae* RNA samples were purified using the RiboPure™ RNA Purification Kit for yeast (Thermo Fisher Scientific Inc., Waltham, MA, USA), following the manufacturer’s instructions. Subsequently, the RNA samples were air-dried in a biosafety hood for 20 hours and preserved in RNA stabilization tubes (GENEWIZ, South Plainfield, NJ, USA) at room temperature before being shipped to Azenta Life Sciences (Suzhou, China) for transcriptome sequencing and analysis. The sequencing was carried out on the Illumina Novaseq platform, using a 2 x 150 bp paired-end (PE) configuration, with approximately 9.0 Gb of PF data per sample. In total, approximately 108.0 Gb of PF data was generated for the 12 *S. cerevisiae* RNA samples, with triplicate samples for each treatment.

### mRNA Library Construction and Sequencing

A total of 1 μg of RNA was used for library preparation. Poly(A) mRNA isolation was performed using Oligo(dT) beads. mRNA fragmentation was achieved using divalent cations and high temperature. Random primers were used for priming. First-strand cDNA and second-strand cDNA were synthesized. The purified double-stranded cDNA was then treated to repair both ends, and dA-tailing was added in a single reaction, followed by a T-A ligation to attach adaptors to both ends. Size selection of adaptor-ligated DNA was performed using DNA Clean Beads. Each sample was amplified by PCR using P5 and P7 primers, and the PCR products were validated.

Libraries with different indexes were multiplexed and loaded onto an Illumina HiSeq, Illumina Novaseq, or MGI2000 instrument for sequencing, using a 2×150 paired-end (PE) configuration following the manufacturer’s instructions.

### Transcriptomic Data Analysis

#### Quality control

To eliminate technical sequences such as adapters, polymerase chain reaction (PCR) primers, or their fragments, and to filter out bases with a quality score lower than 20, the pass filter data in fastq format were processed using Cutadapt (V1.9.1). The parameters used for this processing included a phred cutoff of 20, an error rate of 0.1, an adapter overlap of 1 bp, a minimum length of 75, and a maximum proportion of N of 0.1. This resulted in the generation of high-quality, clean data.

#### Alignment

First, reference genome sequences and gene model annotation files from relevant species were downloaded from genome websites like UCSC, NCBI, and ENSEMBL. Then, the reference genome sequences were indexed using Hisat2 (v2.0.1). Finally, the clean data were aligned to the reference genome using Hisat2 (v2.0.1).

#### Expression analysis

Initially, transcripts in FASTA format were generated from a known GFF annotation file and correctly indexed. Then, using this file as the reference gene file, HTSeq (v0.6.1) estimated gene and isoform expression levels from the paired-end clean data.

#### Differential expression analysis

Differential expression analysis utilized the DESeq2 Bioconductor package, which is based on the negative binomial distribution. The estimates of dispersion and logarithmic fold changes incorporate data-driven prior distributions. Genes with a P-adj value less than 0.05 were considered differentially expressed.

#### GO and KEGG enrichment analysis

GOSeq (v1.34.1) was used to identify Gene Ontology (GO) terms annotating a list of enriched genes with a significant p-adj less than 0.05. Additionally, we utilized topGO to create Directed Acyclic Graphs (DAGs). KEGG (Kyoto Encyclopedia of Genes and Genomes) is a collection of databases encompassing genomes, biological pathways, diseases, drugs, and chemical substances (source: http://en.wikipedia.org/wiki/KEGG). We employed in-house scripts to enrich significantly differentially expressed genes in KEGG pathways.

### Cell Lysates Preparation for LC-MS/MS

*S. cerevisiae* cells were freshly grown in YPD broth overnight. Cell quenching was achieved by transferring 1 mL of cultured broth to a tube containing 4 mL of pre-cooled MeOH/ddH2O (60:40) with 10 mM ammonium acetate. The mixture was promptly placed at -80°C for 2 minutes, followed by centrifugation at 4,000 rpm for 5 minutes at -10°C. Intracellular metabolites were extracted by resuspending the cell pellets in 1 mL of 80% EtOH, followed by heating to 80°C for a total of 4 minutes with 10 seconds of vigorous vortexing in between. Finally, the mixture was centrifuged at 10,000 rpm for 5 minutes at -10°C to obtain the supernatant containing intracellular metabolites. The extracts were aliquoted and stored at - 80°C until analysis using Liquid Chromatography with tandem mass spectrometry (LC-MS/MS), which was conducted by the Proteins & Metabolites Team at AgResearch (Lincoln, New Zealand). The analysis was performed on five biological replicates for each *S. cerevisiae* treatment.

### Metabolomic Analysis

#### Sample preparation

Samples obtained from the cell lysis protocol were stored at -80°C until analysis. To prepare the samples, they were thawed overnight at 4°C, and 800 µL of ice-cold methanol:water (1:1) was added to 200 µL of each sample. The samples were then vigorously shaken using a TissueLyser (Qiagen, USA) and centrifuged for 20 minutes at 4°C (14001 ×g). Two aliquots, each containing 200 µL of the supernatant, were extracted—one for sample analysis and the other for pooled quality control (QC). The samples, along with 200 µL of the pooled QC samples, were dried down using a vacuum concentrator (Vacuumbrand, Wertheim, Germany) and then reconstituted in acetonitrile:water (1:1). D2-tyrosine was added as an internal standard to monitor sample degradation.

#### Chromatography

Samples were analyzed using a Nexera X2 ultra high-performance liquid chromatography (UHPLC) system (Shimadzu, Japan), which included a SIL-30AC autosampler coupled to an LCMS-9030 quadrupole time-of-flight (Q-TOF) mass spectrometer (Shimadzu, Japan) equipped with an electrospray ionization source (see Figure S11). A 2 µL sample was injected into a normal phase Ascentis® Express HILIC UHPLC column (2.1 x 100 mm, 2 µm particle size; Sigma, USA) and eluted at 30°C over a 20-minute gradient with a flow rate of 400 µL/min. The mobile phase consisted of solvent A (10 mM ammonium formate in water) and solvent B (acetonitrile with 0.1% formic acid). The solvent gradient program started at 97% solvent B from 0 to 0.5 minutes, decreased to 70% within 11.5 minutes, further decreased to 10% from 11.5 to 13.5 minutes, held at 10% for 1.5 minutes, increased to 97% B within 1 minute, and remained at that concentration until the end of the elution run.

#### Mass spectrometry

Full scans (m/z 55-1100) and MS/MS scans for windows spanning m/z 20 were set up for analyses in both positive and negative ionization modes. A total of 42 events were configured with a loop time of 0.85 seconds. The spray voltage was set at 4.0 kV for positive ionization mode and -3.0 kV for negative ionization mode. The collision energy (CE) was maintained at 23 ± 15 V. The ion source was operated under optimal conditions, including a nebulizing gas flow of 3.0 L/min, a heating gas flow of 10.0 L/min, an interface temperature of 300°C, a drying gas flow of 10.0 L/min, a desolvation line temperature of 250°C, and a heat block temperature of 400°C.

#### Batch sequence

All samples were analyzed in a single batch, beginning with positive ionization mode. The sequence started with five blanks, followed by an external standard (Amino acid standard, A9906, Sigma, USA) to verify system performance. Next, a few QC pooled samples were run, and then the actual samples in a randomized order with QC samples interspersed once every 8 samples. The same order was followed for negative ionization mode.

#### Data analysis

Raw data files (.lcd) were converted to the mzML file format using LabSolutions software (Version 5.99 SP2, Shimadzu, Japan). MS-Dial was employed for peak detection, MS2 deconvolution, sample alignment, and compound identification^91^. Appropriate adducts were selected for both positive and negative ionization modes. Blank data was subtracted from all samples, ensuring that the sample’s maximum ratio to the average of the blanks was maintained at 5. Local Weighted Scatterplot Smoothing (LOWESS) was used for signal correction during data preprocessing. After preprocessing, only features with metabolite IDs that matched with MS/MS or m/z (without MS/MS) were exported, along with their respective normalized peak areas for statistical analysis. Statistical analysis of the normalized data was performed using MetaboAnalyst 5.0, a web-based platform for metabolomics data analysis^92^.

### Yeast Viability and Morphology

Overnight *S. cerevisiae* cultures were grown in YPD broth, inoculated into flasks with a flask volume to medium volume ratio of 5:1, and incubated at 32°C with shaking at 120 rpm. Yeast viability was assessed using both optical density at 600 nm and colony-forming units (CFU). Optical density was measured with a FLUOstar Omega microplate reader (BMG LABTECH, Ortenberg, Germany) at relevant dilution factors. CFU counts were obtained by spotting 10 µL of serially diluted cultured broth onto YPD plates, which were then counted after 24 hours of incubation at 37°C. The results were reported as the viability (CFU/mL) for each strain. The data represents the average of two independent experiments conducted simultaneously, with technical triplicates for each experiment. A 1X phosphate-buffered saline (PBS) solution was used as the diluent for serial dilution. Yeast morphology was observed and recorded using a Gel Doc XR+ Gel Documentation System (Bio-Rad Laboratories, Hercules, CA, USA).

### Gene Phylogeny

The protein sequences of selected upregulated genes associated with the autophagy process (SSE1, NCBI accession NP_015219.1) and stress response (TOR1, NCBI accession NP_012600.1; SKN1, NCBI accession NP_011659.3), as well as acetyltransferase (EHT1, NCBI accession NP_009736.3) and methyltransferase (COQ5, NCBI accession NP_013597.1), were searched against the Swiss-Prot database using PROSITE with an E-value threshold of 10^-5^ ^93^. Additional homologs were identified using the curated databases on SHOOT.bio^94^ and the ‘top IMG homologs’ function on the IMG website^95^. Further homologs were obtained by searching the protein sequences of individual genes against the NCBI non-redundant protein database (nr50) using HMMER (v. 3.3)^96^ within the MPI bioinformatics toolkit^97^, with an e-value threshold of 10^-5^ and retaining the default values of other parameters.

The sequences of homologs obtained for each gene were clustered with DIAMOND-DeepClust (v. v2.1.3.157)^98,99^ within the MPI bioinformatics toolkit, with default parameters. Whenever possible, gene sequences from the Asgard group were used as outgroups, as suggested elsewhere^100^. The query sequences of individual genes, clustered homologous sequences and outgroup sequences were then aligned using MAFFT (v. 7.273)^101^, with a gap opening penalty of 3^102^. The resulting alignment was used as input for phylogenetic tree construction with IQ-TREE (v. 1.6.12)^103^. The best phylogenetic model was determined using ModelFinder^104^ within IQ-TREE. The reliability of the nodes in the tree was estimated with 1000 ultrafast bootstraps^105^ and a maximum of 3000 iterations. Finally, the phylogenetic tree was visualized and annotated using iTOL (v.6.8)^106^.

### Gene Co-occurrence Pattern

The protein sequences of genes related to the autophagy process, stress response, lifespan/aging, acetyltransferases, and methyltransferases in *S. cerevisiae* S288C were downloaded from the NCBI GenBank database. These gene protein sequences were then input into STRING^107^ (https://string-db.org/) using default parameters to visualize their co-occurrence across different domains of life.

### Data Visualization and Statistical Analysis

Microsoft Office (Microsoft Corporation, Redmond, WA, USA) was used for creating basic graphs. RStudio (Posit, Boston, MA, USA) was employed for generating transcriptomic heatmaps with DESeq2, ggplot2, and ComplexHeatmap packages, as well as multi-omics graphs using the mixOmics package. Circos version 0.69-9 was utilized for visualizing metabolomics data. PCA and AHC were analyzed with XLSTAT Statistical Software 2016 (Addinsoft, Paris, France). The statistical analysis was performed with a 95% confidence level, and the data were presented as mean ± SD.

## Author Contributions

V.C. conceived the study; Y.K., D.A., and A.S (1). performed research; Y.K., C.W., S.L.W.O., A.S. (1), A.S. (2), V.C., analyzed and interpreted data, Y.K., C.W., S.L.W.O, P.A.W. and V.C. wrote the paper.

## Acknowledgments

Callaghan Innovation R&D Fellowship Grant (Grant No: 70570) and KiwiNet PSAF (Grant No: 46561) funded this work. We also extend our thanks to Dr. Nadia Mitchell and Dr. Samantha Murray, for their valuable contributions.

