## Supplementary material for "Reevaluating the Concept of Aging: Long-Term Stress Adaptation as a Key Factor in Yeast Aging": Supplementary File - 03_11_23.pdf

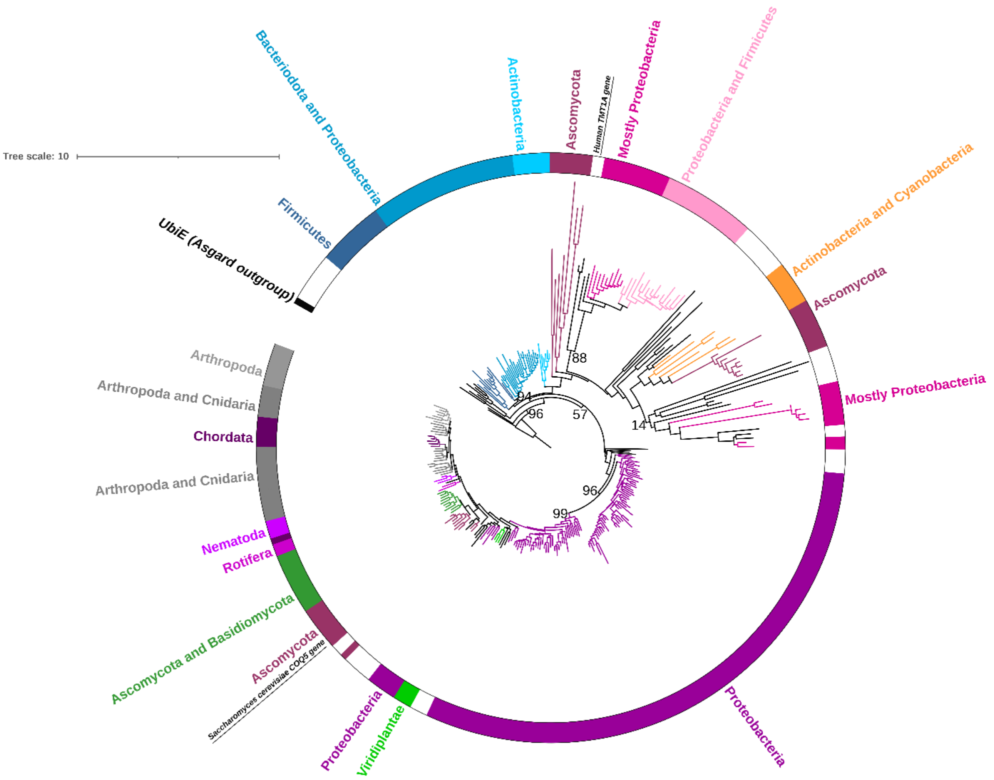


**Supplementary Figure 1** Maximum likelihood phylogenetic tree of the *S. cerevisiae coq5* gene and its homologs across different domains of life. The phylogenetic model employed was LG+I+G4, while numbers are ultrafast bootstrap values for selected branches. The branches are coloured according to the main phyla/groups. The outer ring is coloured according to the main phyla/groups, while branches with dissimilar members are uncoloured.

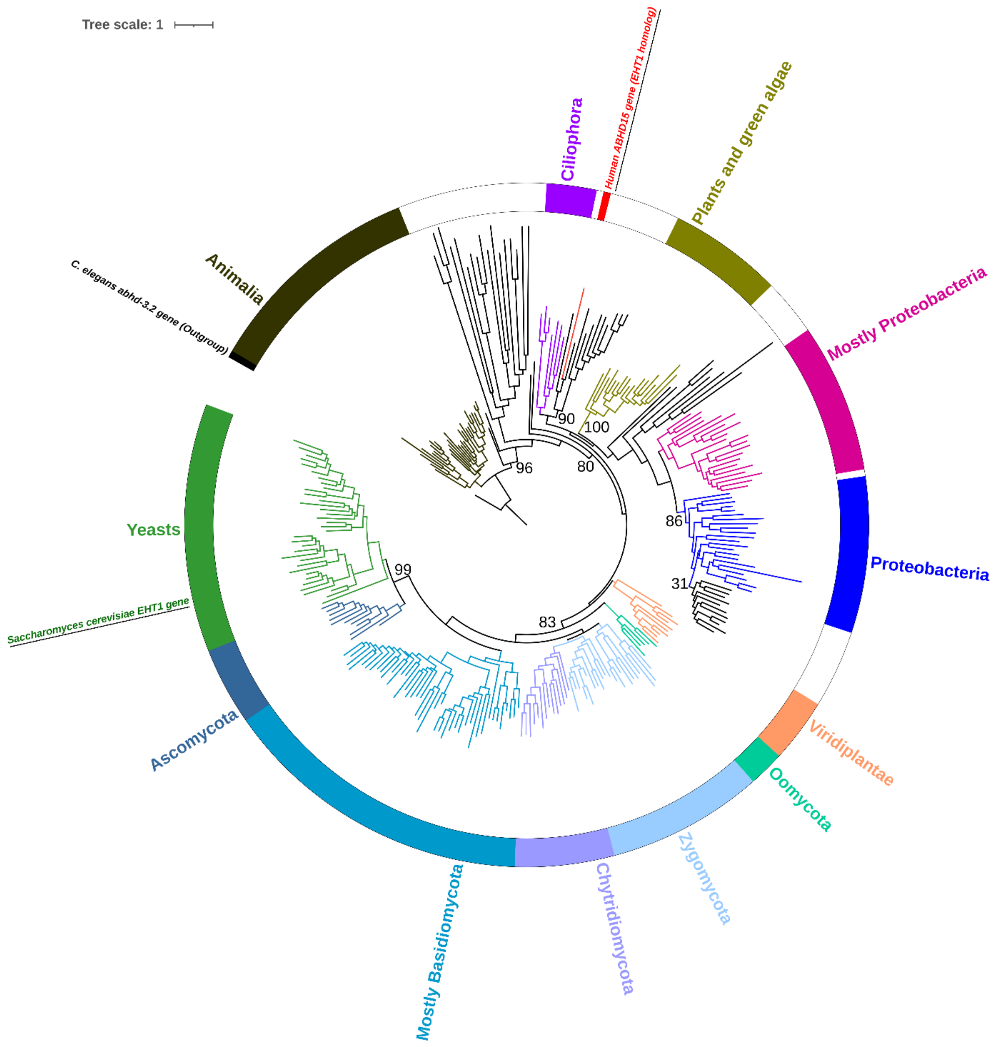


**Supplementary Figure 2** Maximum likelihood phylogenetic tree of the *S. cerevisiae eht1* gene and its homologs across different domains of life. The phylogenetic model employed was LG+G4, while numbers are ultrafast bootstrap values for selected branches. The branches are coloured according to the main phyla/groups. The outer ring is coloured according to the main phyla/groups, while branches with dissimilar members are uncoloured.

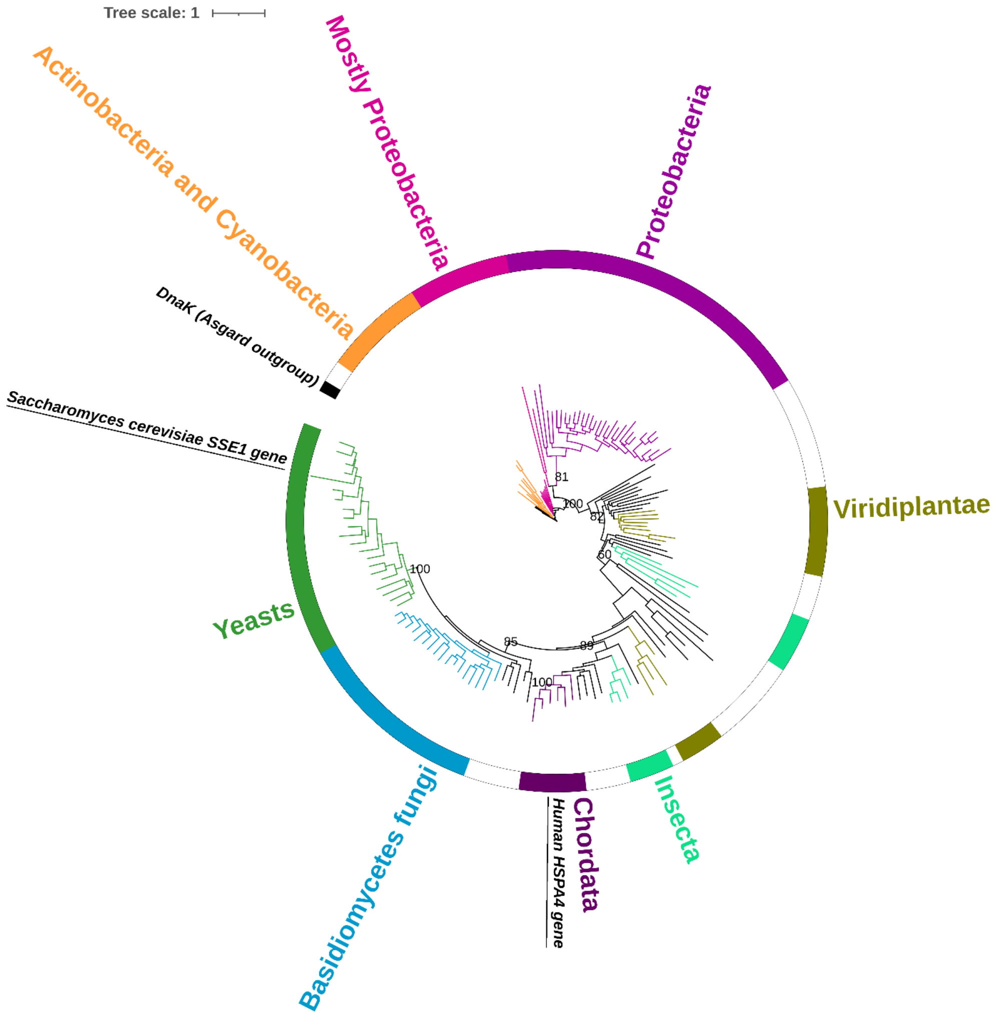


**Supplementary Figure 3** Maximum likelihood phylogenetic tree of the *S. cerevisiae sse1* gene and its homologs across different domains of life. The phylogenetic model employed was LG+I+G4, while numbers are ultrafast bootstrap values for selected branches. The branches are coloured according to the main phyla/groups. The outer ring is coloured according to the main phyla/groups, while branches with dissimilar members are uncoloured.

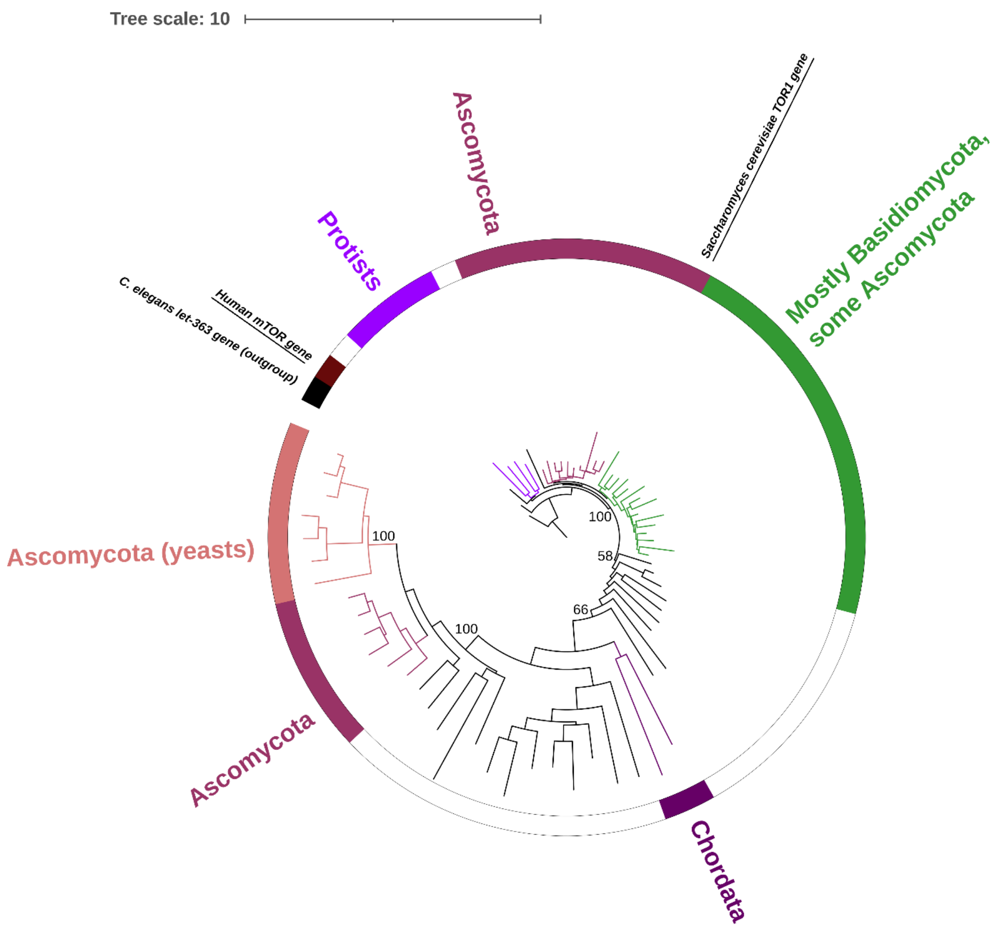


**Supplementary Figure 4** Maximum likelihood phylogenetic tree of the *S. cerevisiae tor1* gene and its homologs across different domains of life. The phylogenetic model employed was LG+F+I+G4, while numbers are ultrafast bootstrap values for selected branches. The branches are coloured according to the main phyla/groups. The outer ring is coloured according to the main phyla/groups, while branches with dissimilar members are uncoloured.

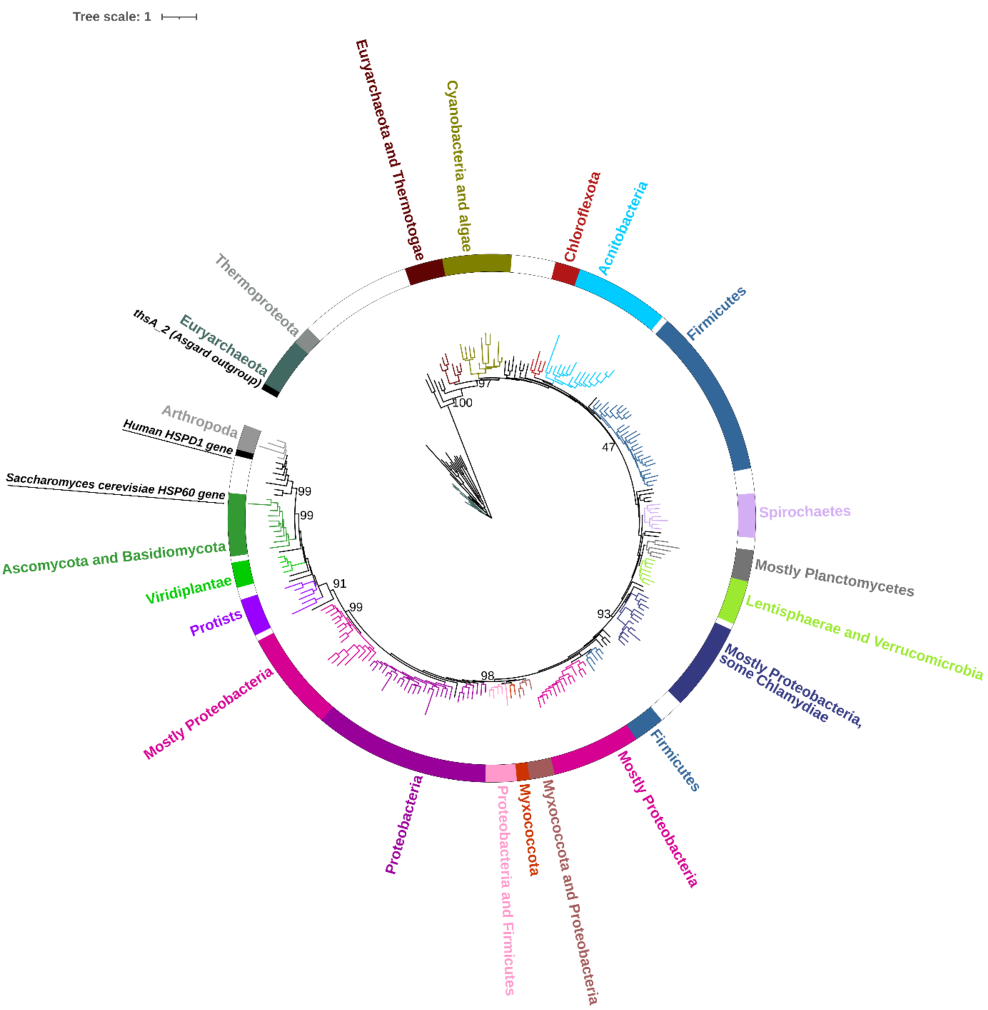


**Supplementary Figure 5** Maximum likelihood phylogenetic tree of the *S. cerevisiae hsp60* gene and its homologs across different domains of life. The phylogenetic model employed was LG+F+I+G4, while numbers are ultrafast bootstrap values for selected branches. The branches are coloured according to the main phyla/groups. The outer ring is coloured according to the main phyla/groups, while branches with dissimilar members are uncoloured.

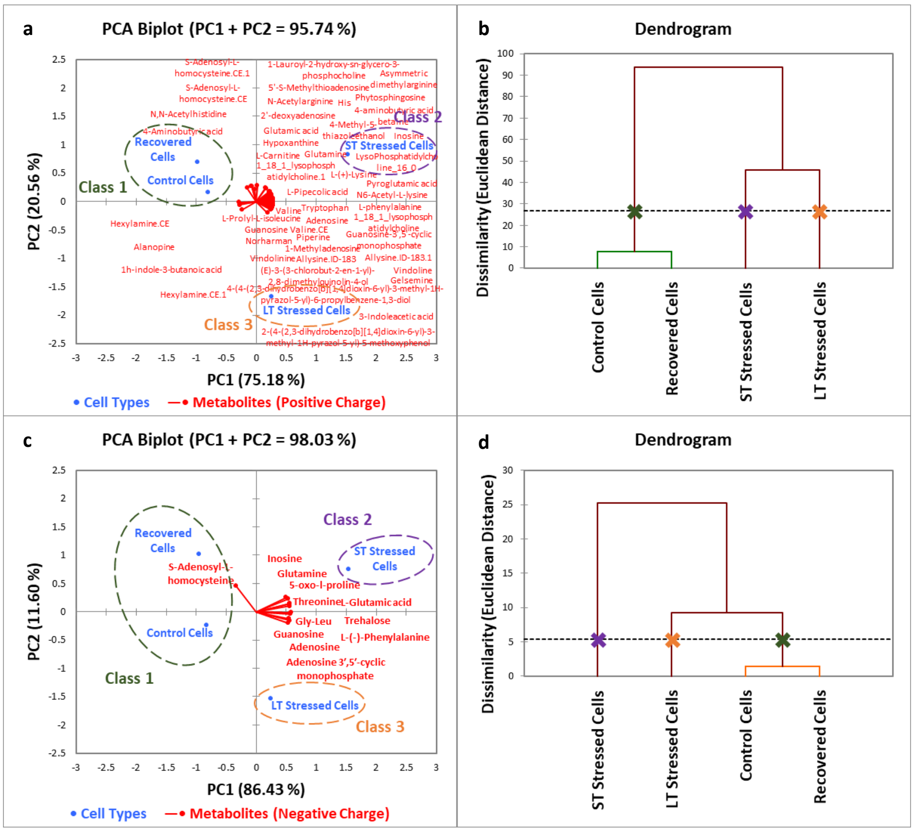


**Supplementary Figure 6 a.** Principal component analysis (PCA) biplot illustrating the relationship between *S. cerevisiae* cell types and known metabolites detected by LC-MS/MS analysis under positive charge mode. **b.** *S. cerevisiae* cell types grouped using a agglomerative hierarchical clustering (HCA) according to their dissimilarity levels based on metabolites detected under positive charge mode. **c.** Principal component analysis (PCA) biplot illustrating the relationship between *S. cerevisiae* cell types and known metabolites detected by LC-MS/MS analysis under negative charge mode. **d.** *S. cerevisiae* cell types grouped using a agglomerative hierarchical clustering (HCA) according to their dissimilarity levels based on metabolites detected under negative charge mode.

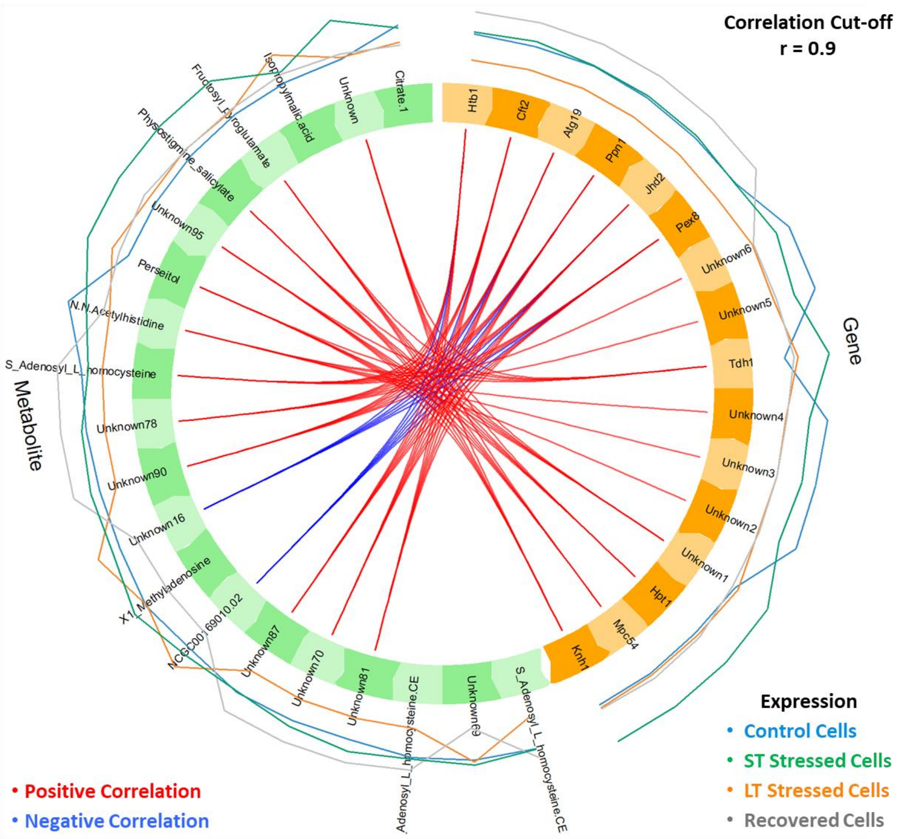


**Supplementary Figure 7** Circos plot from multiblock sPLS-DA performed on the *S. cerevisiae* RNA sequencing and LC-MS/MS metabolomics data (by Mixomics package in R). The plot represents the correlations greater than 0.9 between variables of different types, represented on the side quadrants. The internal connecting lines show the positive and negative correlations. The outer lines show the expression levels of each variable (gene and metabolite) in each yeast cell type.

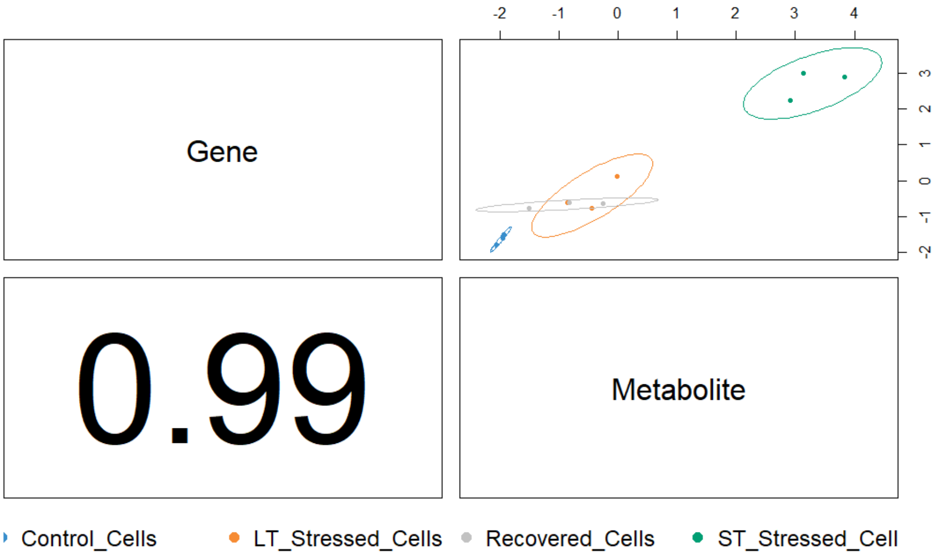

**Supplementary Figure 8** Diagnostic plot from multiblock sPLS-DA analysis grouping treated and untreated *S. cerevisiae* in terms of the selected genes and metabolites (ncomp. = 3).

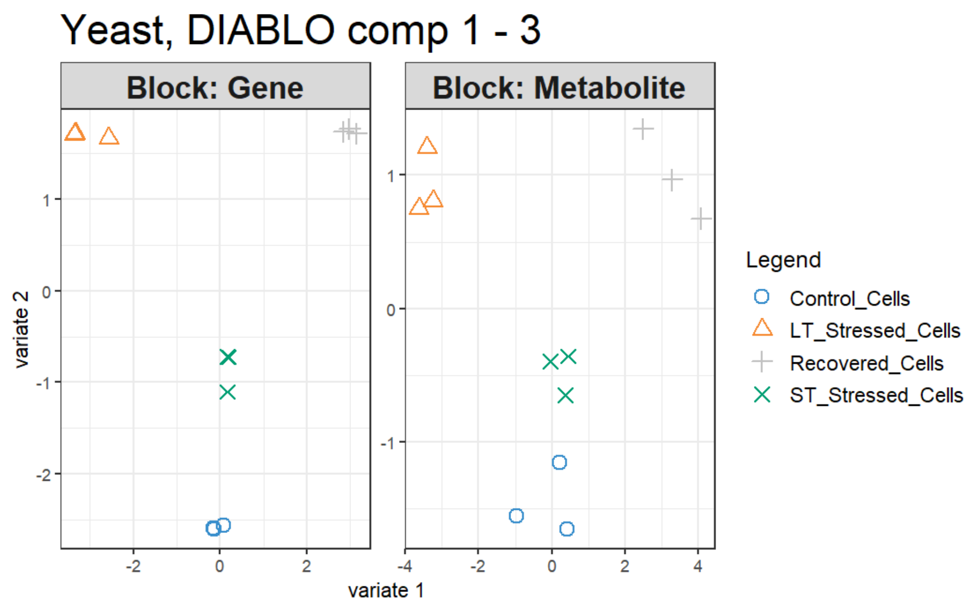


**Supplementary Figure 9** *S. cerevisiae* samples plot by multiblock sPLS-DA analysis, the samples are plotted according to their scores on the first 2 components for each data set. The plot shows the degree of agreement between the different data sets and the discriminative ability of each data set.

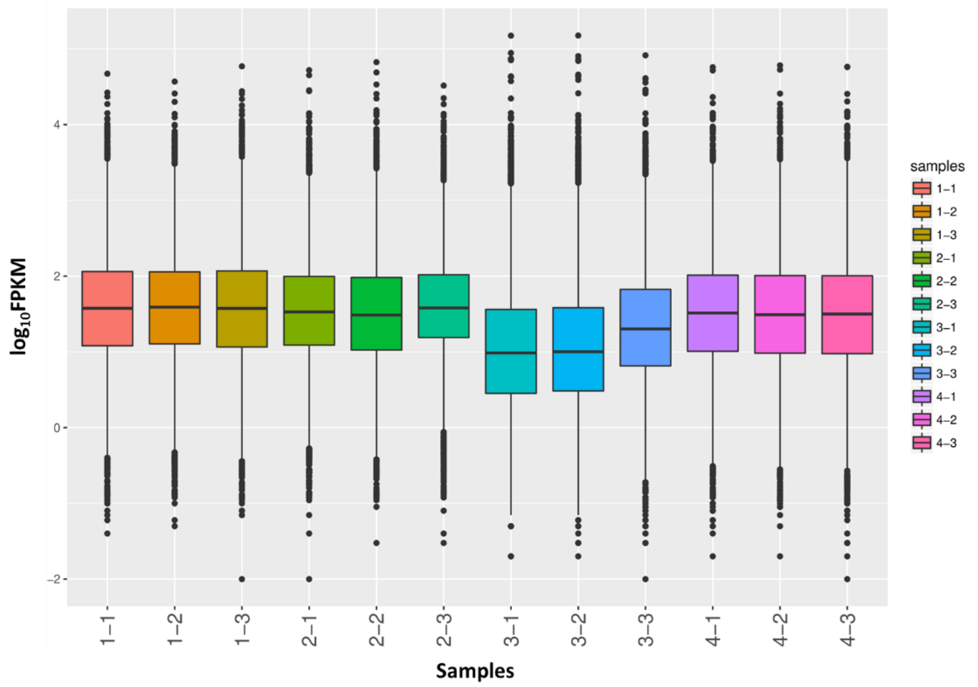


**Supplementary Figure 10** Comparison of gene expression levels (logFPKM) under different experimental conditions. Each of the five elements in each box plot, from top to bottom, specifies the maximum, upper quartile, median, lower quartile and the minimum value, respectively.

**1-1, 1-2 & 1-3: Control Cells,** wild type *S. cerevisiae*; **2-1, 2-2 & 2-3: ST Stressed Cells,** short-term stressed cells, *S. cerevisiae* firstly exposed to 10 mM benzoic acid for 16 hrs; **3-1, 3-2 & 3-3: LT**

**Stressed Cells,** long-term stressed cells, *S. cerevisiae* exposed to 10 mM benzoic acid for 500 hrs (24 hrs/sub-culture); **4-1, 4-2 & 4-3: Recovered Cells,** *S. cerevisiae* exposed to 10 mM benzoic acid for 500 hrs (24 hrs/sub-culture), followed by growing in regular YPD broth for 16 hrs.

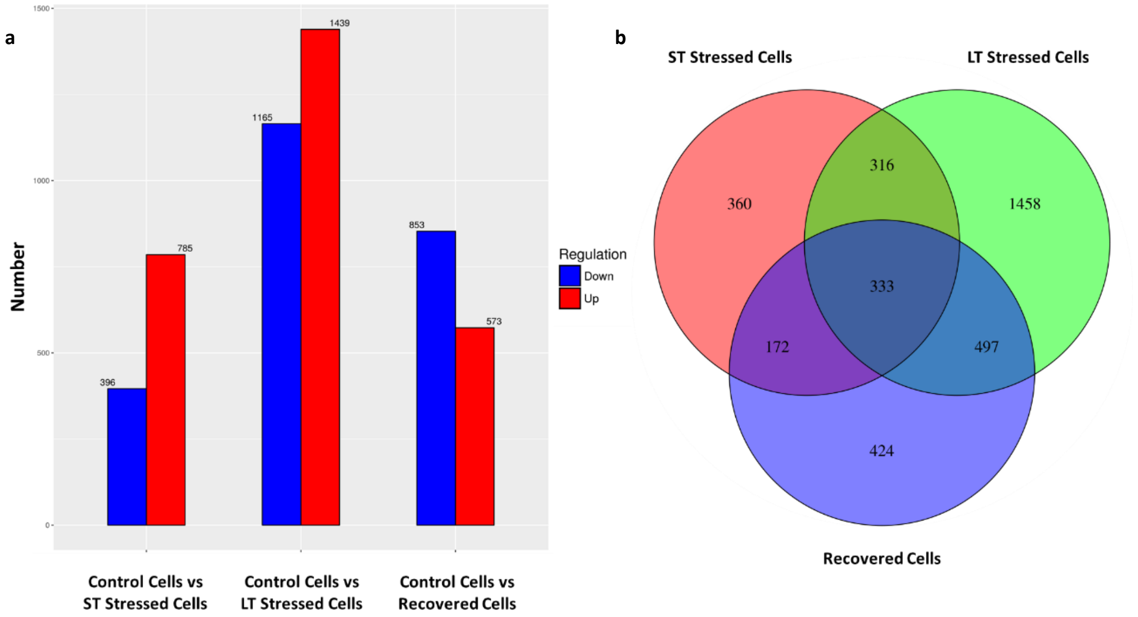


**Supplementary Figure 11 a.** Bar graph showing the number of genes significantly up- or down-regulated in treated groups compared to the control. **b.** Venn diagram illustrating the relationships among all treated groups compared to the control.

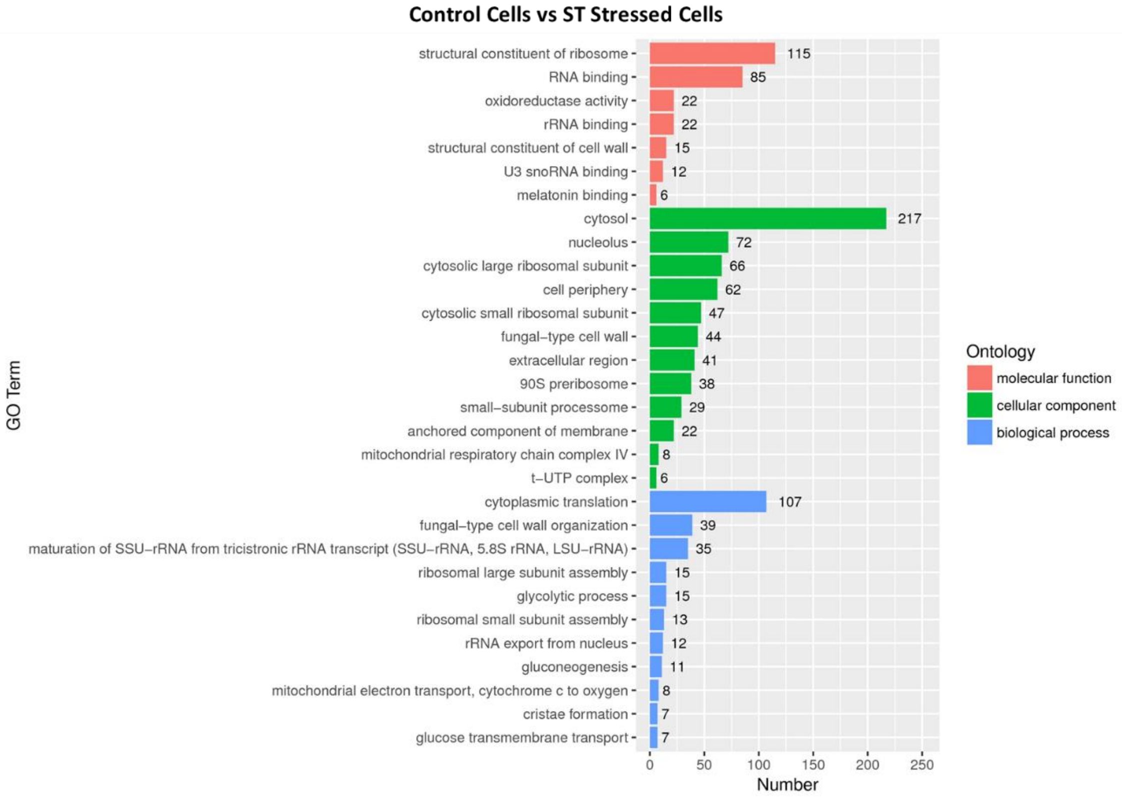

**Supplementary Figure 12** Histogram depicting GO enrichment analysis of short-term (ST) stressed cells in comparison to control cells. The X-axis represents the number of differentially expressed genes within each GO category. Different categories, such as biological processes, cellular components, and molecular functions, are distinguished by colour codes on the right side.

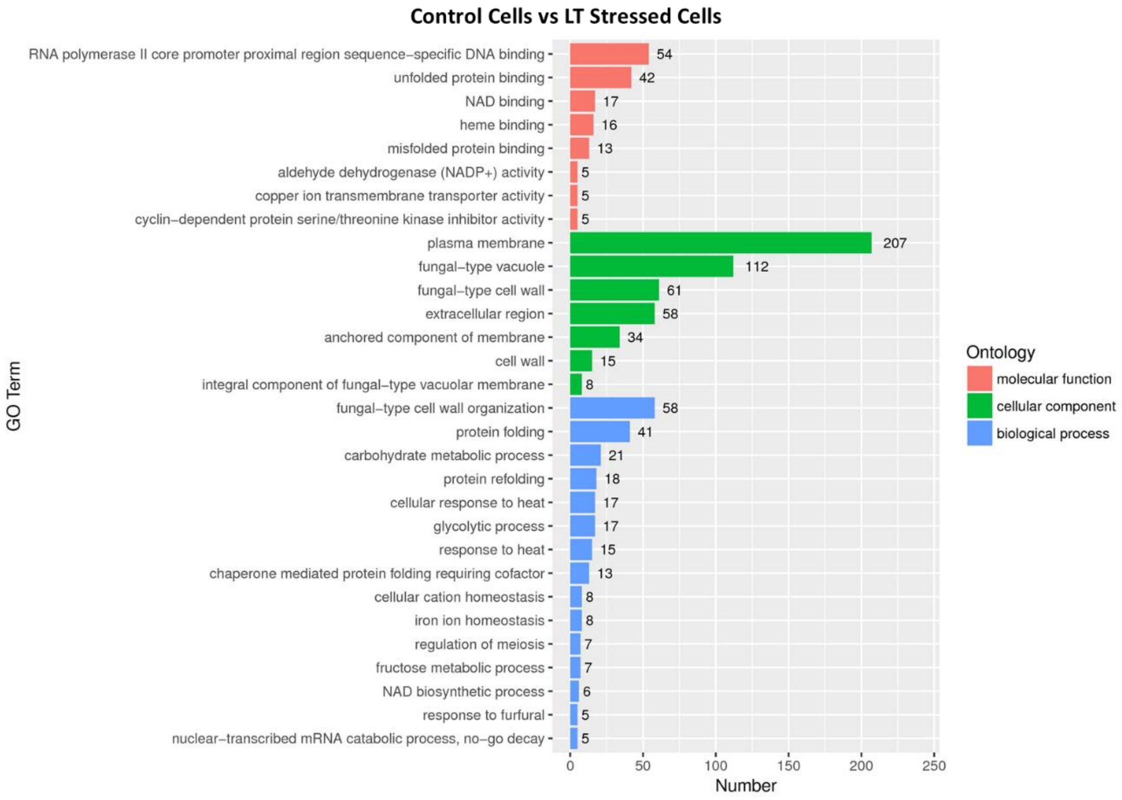


**Supplementary Figure 13** Histogram depicting GO enrichment analysis of long-term (LT) stressed cells in comparison to control cells. The X-axis represents the number of differentially expressed genes within each GO category. Different categories, such as biological processes, cellular components, and molecular functions, are distinguished by colour codes on the right side.

### Control Cells vs Recovered Cells

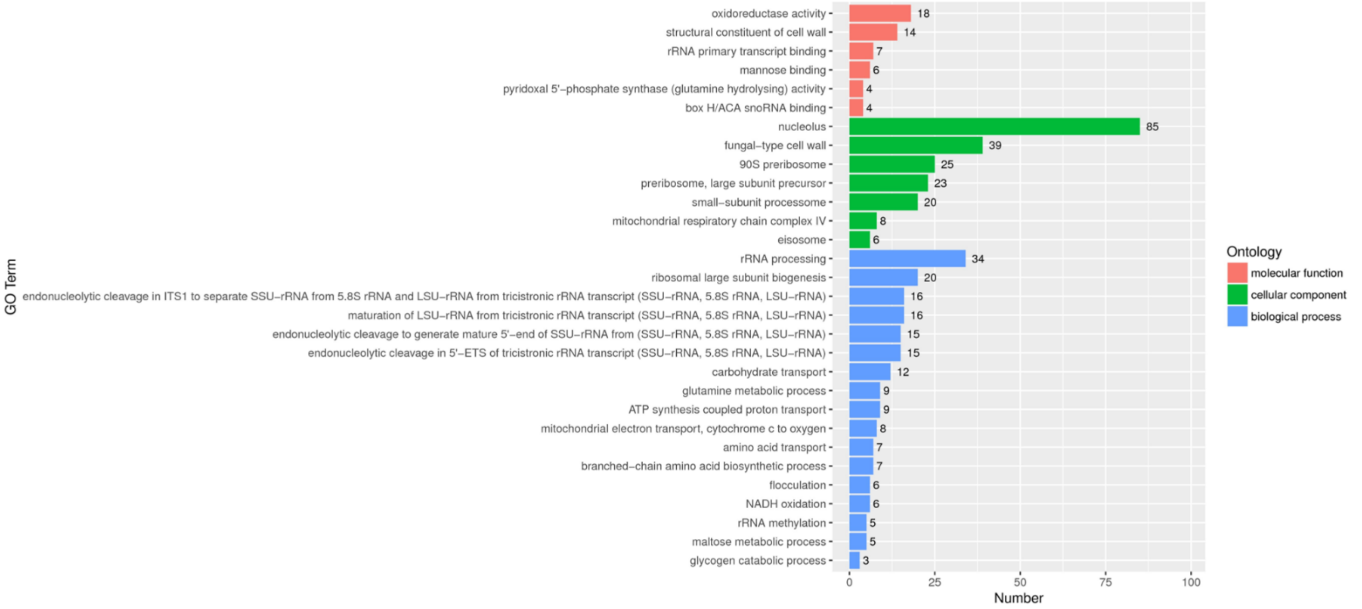


**Supplementary Figure 14** Histogram depicting GO enrichment analysis of recovered cells in comparison to control cells. The X-axis represents the number of differentially expressed genes within each GO category. Different categories, such as biological processes, cellular components, and molecular functions, are distinguished by colour codes on the right side.

### Control Cells vs ST Stressed Cells

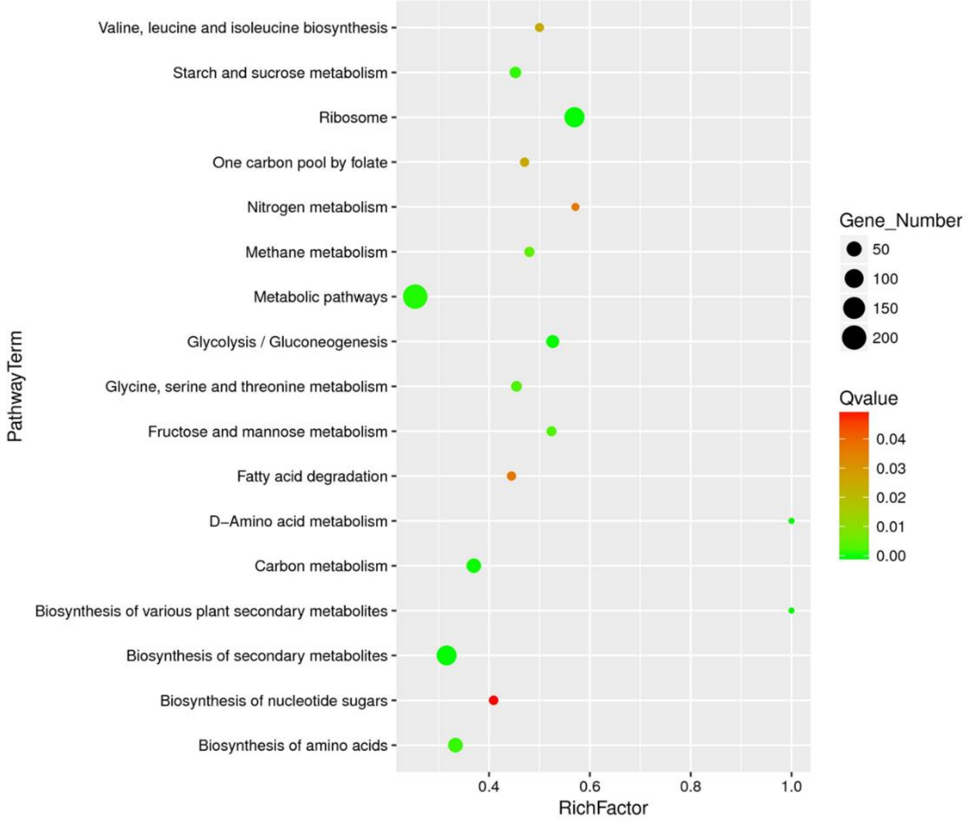


**Supplementary Figure 15** Scatter plot depicting KEGG enrichment of differentially expressed genes in short-term (ST) stressed cells compared to the control cells. Dot size corresponds to the number of differential genes within each pathway, showing a positive correlation. Different Q-value ranges are represented by distinct colours.

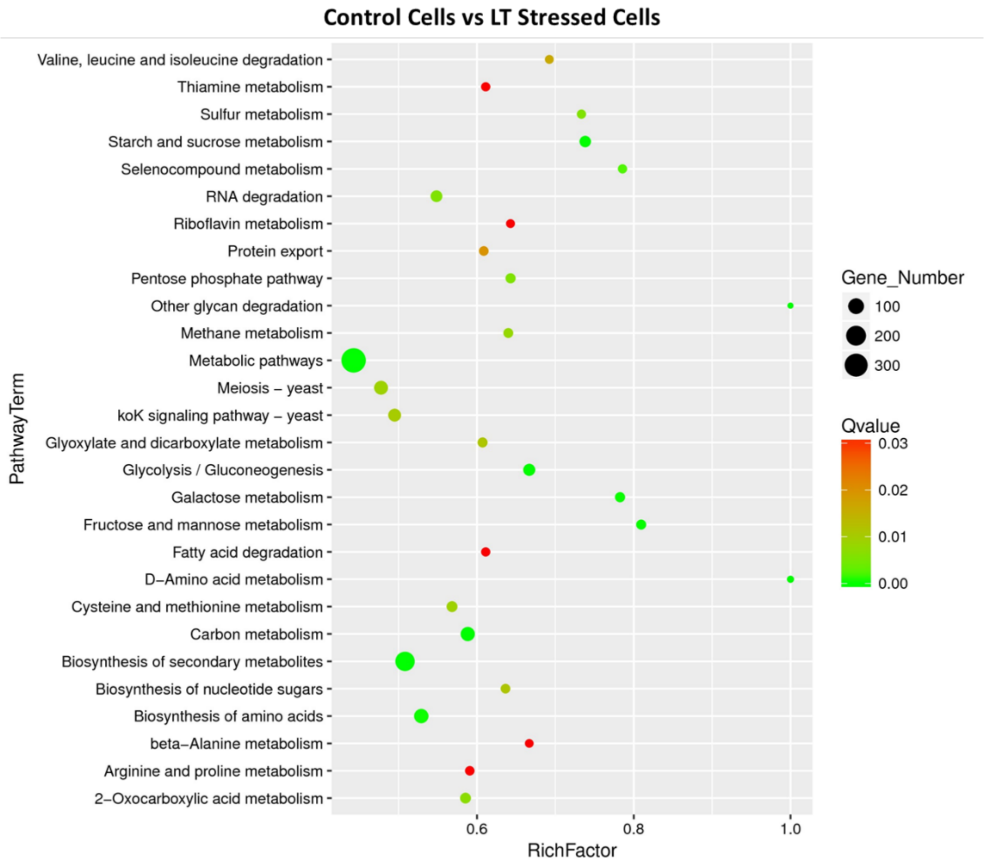


**Supplementary Figure 16** Scatter plot depicting KEGG enrichment of differentially expressed genes in long-term (LT) stressed cells compared to the control cells. Dot size corresponds to the number of differential genes within each pathway, showing a positive correlation. Different Q-value ranges are represented by distinct colours.

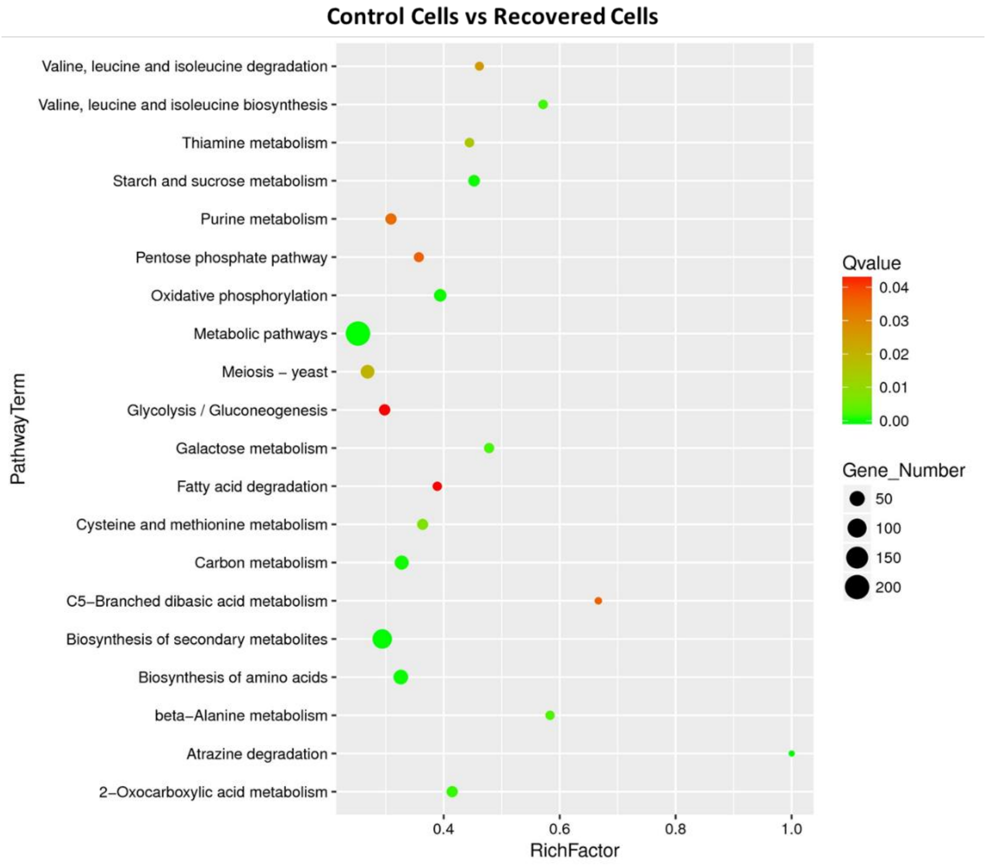


**Supplementary Figure 17** Scatter plot depicting KEGG enrichment of differentially expressed genes in recovered cells compared to the control cells. Dot size corresponds to the number of differential genes within each pathway, showing a positive correlation. Different Q-value ranges are represented by distinct colours.
